# Smarcc1 is essential for the patterning of the optic stalk and differentiation of the optic nerve head astrocytes

**DOI:** 10.64898/2025.12.30.697056

**Authors:** Nitay Zuk-Bar, Shai Ovadia, Guizhong Cui, Alexey Obolensky, Eyal Banin, Ron Ofri, Naihe Jing, Ruth Ashery-Padan

## Abstract

The optic nerve is essential for vision, and its development depends on the transient neuroectodermal optic stalk, which forms alongside the optic cup through coordinated morphogenesis and differentiation into optic nerve astrocytes that provide lifelong support for retinal ganglion cell (RGC) axons. Here, we delineate key steps in astrocyte formation from the optic stalk and uncover stage- and lineage-specific roles of Smarcc1 (Baf155) and Smarcc2 (Baf170), scaffolding subunits of SWI/SNF chromatin remodeling complexes. Both factors are co-expressed in early retinal pigment epithelium (RPE) and optic stalk progenitors, with Smarcc2 persisting in differentiated derivatives, including RPE and optic nerve astrocytes. Conditional deletion using Dct-Cre revealed reciprocal compensation in pigmented lineages, whereas Smarcc1 loss uniquely caused optic nerve head (ONH) morphogenetic failure, characterized by glial lamina collapse, RGC degeneration, and progressive visual decline. Spatial transcriptomics and functional assays demonstrate that Smarcc1 drives the transition of dorsal optic stalk progenitors from a pigmented, RPE-like state to astrocyte progenitors by repressing pigment gene programs and enabling Pax2/Sox2 activity. After specification, Smarcc1 is required for glial lamina assembly, essential for ONH integrity and RGC survival. These findings establish a mechanism in which Smarcc1-mediated chromatin remodeling couples astrocyte progenitor specification and differentiation with local morphogenesis, ensuring long-term retinal function.

## Introduction

Complex interactions between transcription factors and chromatin remodeling complexes govern the stage- and cell-specific gene expression programs required for organ formation. In the central nervous system (CNS), this interplay is exemplified by the transition of neuroectodermal progenitors into diverse neural and glial lineages. A striking case is the optic nerve, which arises from the transient neuroepithelial optic stalk (OS). Initially serving as a conduit for retinal ganglion cell (RGC) axons extending to the brain, the OS later differentiates into astrocytes that support and ensheathe these axons. Here, we delineate the spatial and temporal dynamics underlying the conversion of the optic stalk into optic nerve astrocytes. Our findings reveal partial redundancy between Smarcc1 (Baf155) and Smarcc2 (Baf170), core subunits of the SWI/SNF chromatin remodeling complex, and uncover a specific, stage-dependent requirement for Smarcc1 in the specification and differentiation of optic nerve astrocytes.

The optic nerve transmits visual information through RGC axons and is supported by astrocytes and a vascular network (Butt et al., 2004). At its anterior end, the optic nerve head (ONH) serves as the convergence point of all RGC axons. In both humans and mice, axons remain unmyelinated within the ONH but acquire myelin posteriorly. The ONH comprises the optic disc within the eye cup and the optic nerve laminar region (ONLR) just behind it. (Bernstein et al., 2020; Morcillo et al., 2006; Sun et al., 2009). The ONH also contains Sox2-positive neural precursors that contribute to postnatal optic nerve extension (Bernstein et al., 2020). The optic nerve head integrity is crucial for RGC survival throughout life, and structural changes in this region are linked to glaucomatous optic neuropathy (Howell et al., 2007; Pitha et al., 2024).

During embryogenesis, the optic stalk originates from the proximal optic vesicle, a neuroectodermal outgrowth of the forebrain (Colello and Guillery, 1992). The distal vesicle invaginates to form the optic cup, which consists of an outer layer of the progenitors for the presumptive retinal pigmented epithelium (pRPE) and an inner layer of retinal progenitor cells (RPCs). The proximal region of the optic vesicle narrows into the epithelial tube of the optic stalk. A ventral fissure allows entry of hyaloid vasculature and mesenchymal cells (Barishak, 2001; Lutty and McLeod, 2018; Saint-Geniez and D’Amore, 2004), which ultimately close as the fissure fuses to form a continuous tube ensheathing emerging RGC axons (Silver and Robb, 1979). Over developmental time, optic stalk progenitors delaminate and differentiate into type I astrocytes (Cardozo et al., 2020; Heavner and Pevny, 2012; Tao and Zhang, 2014). Yet, how regional identity is established within the optic stalk, and how progenitors are directed toward an astroglial fate, are still unresolved questions.

Patterning of the optic stalk and optic cup arises from positional cues—Shh at the midline, Tgfb from mesenchyme, and Fgfs/WNTs from surface ectoderm (Cardozo et al., 2020; Morcillo et al., 2006). These factors instruct the expression of a combination of region-specific transcription factors, which in turn trigger the gene regulatory networks required for regional cell fates and, at the same time, inhibit competing developmental programs (Cardozo et al., 2020; Patel and Sowden, 2019). An example of such TF interplay is the formation of optic stalk and optic cup progenitor territories through inhibitory interactions between the paired type homeodomain factors, Pax2 and Pax6. Notably, Pax2 and Pax6 act in mutual opposition: Pax2 marks optic stalk progenitors and restricts them to astroglial lineages, whereas Pax6 promotes optic cup fates. Loss of Pax2 disrupts the stalk–cup boundary, causes Pax6 upregulation in stalk cells, and prevents optic fissure closure, resulting in coloboma (Bosze et al., 2021; Schwarz et al., 2000). Although transcription factor interplay has been well studied in this context, much less is known about how chromatin remodeling complexes contribute to stalk patterning and astrocyte specification.

The SWI/SNF (BAF) complexes are critical regulators of neural development, promoting chromatin accessibility in opposition to Polycomb-mediated repression (Ho and Crabtree, 2010; Kadoch et al., 2017). Their core subunits—Smarca4/2 (Brg1/Brm), Smarcb1 (Baf47), and Smarcc1/2 (Baf155/170)—interact with lineage-restricted transcription factors to direct fate specification (Hoffman et al., 2018; Tuoc et al., 2013b). Smarcc1 and Smarcc2 act as interchangeable scaffold components but also exhibit non-redundant developmental functions: Smarcc1 is essential for ESC pluripotency (Ho et al., 2009), and both subunits modulate cortical development by altering Pax6 transcriptional activity (Tuoc et al., 2013a). SWI/SNF activity has also been linked to Sox2 and MITF function in diverse settings (Ahmed et al., 2012; Laurette et al., 2015), and conditional removal of both Smarcc1/2 in the RPE redirects cells to neural-like states (Ovadia et al., 2023). Smarcc1 and Smarcc2 are implicated in a group of neurodevelopmental disorders collectively termed BAF-opathies (Chen et al., 2022), underscoring the importance of dissecting their individual roles, compensatory mechanisms, and functional specificity.

Despite these insights, how Smarcc1 and Smarcc2 coordinate with transcription factors to establish OS identity and promote astrocyte differentiation remains unresolved. Our study addresses this gap by uncovering a temporally restricted requirement for Smarcc1 in optic nerve astrocyte specification, revealing new principles of chromatin–transcription factor interplay in CNS development.

## Results

### Distinct stage and cell-specific expression of Smarcc1 and Smarcc2 in the optic vesicle lineages

To investigate the stage and tissue-specific roles of Smarcc1 and Smarcc2 in the pigmented lineages of the optic cup, we characterized, by antibody labeling, their expression patterns in the neuroectodermal derivatives of the embryonic optic cup.

The two subunits presented a partly overlapping pattern of expression in the optic cup derivatives. At E14 and P0 both subunits were detected in the cells of the OS and in the RPE progenitors (Fig.1A-D). Within the developing retina, at E14 and P0, Smarcc1 was detected in the retinal progenitor cells in the neuroblastic (NBL) layer, while Smarcc2 was mostly in the inner nuclear layer (INL), which is populated by the post-mitotic retinal precursors of ganglion, amacrine, and horizontal retinal cells (Fig. 1A-D). Both subunits were detected in the ciliary body (CB) and iris progenitors at the distal tips of the optic cup (Fig. S1).

**Fig. 1.**
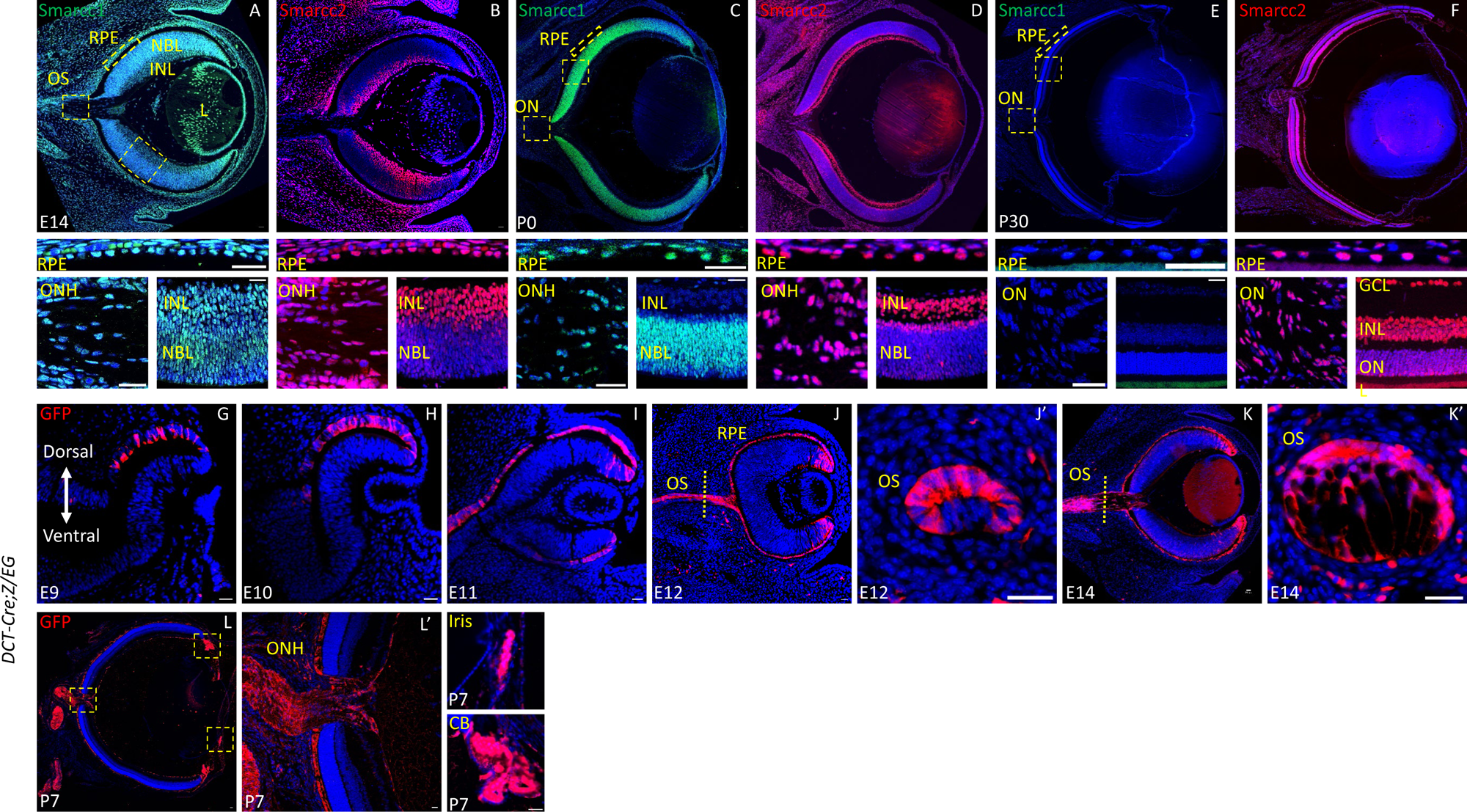
The stage-specific expression of Smarcc1 and Smarcc2 in the optic vesicle lineages and characterization of the *DCT-Cre* activity. Antibody labeling of E14 (A-B), P0 (C-D) and P30 (E-F) for detection of Smarcc1 (A,C,E) and Smarcc2 (B, D, F). Lower panels are the magnifications of the RPE, ONH and retina. The recombination pattern mediated by *DCT-Cre* monitored by antibody labeling for GFP (G-L) expressed by the *Z/EG* reporter at E9 (G), E10 (H), E11 (I), E12 (J), E14 (K) and P7 (L). Higher magnifications of the boxed regions in L are presented on the right (L’). Transverse sections of the OS are shown in J’ and K’. Abbreviations: OS-Optic Stalk, RPE-Retinal Pigmented Epithelium, NBL-Neuroblast Layer, INL-Inner Nuclear Layer, L-Lens, ON-Optic Nerve, ONH-Optic Nerve Head, CB-Ciliary Body. Scale bars = 25µm.

In the mature retina (P30), the expression of Smarcc1 was reduced below detection in the optic cup/stalk lineages, whereas Smarcc2 was detected in the mature cell types that are derived from the optic vesicle: optic nerve astrocytes, RPE, the retinal cell types, ciliary body and iris (Fig. 1E, F, Fig. S1).

The detection of Smarcc2 but not Smarcc1 in differentiated derivatives of the optic cup corresponds with the stage-specific role attributed to each of these subunits; Smarcc1 in proliferating multipotent progenitors and Smarcc2 in differentiating and mature cell types (Bachmann et al., 2016; Bi-Lin et al., 2021; Zhang et al., 2021).

### Conditional deletion of Smarcc1 or Smarcc2 in the pigmented lineages of the eye using *Dct-Cre* includes the RPE and OS

To examine the Smarcc1 and Smarcc2 specific roles and their possible functional redundancy in the developing and mature pigmented lineages of the eye, we selectively deleted one subunit at a time using the *Dct-Cre* (Davis et al., 2009). To trace the trajectory of Cre-expressing cells, we included the *Z/EG* reporter in which eGFP is activated in cells where Cre recombinase was active (Novak et al., 2000) (Fig. 1G-L). The detailed analysis of the eGFP in the *Dct-Cre;Z/EG* embryos revealed that, in addition to the documented activity of the *Dct-Cre* in pRPE in the dorsal optic cup (Davis et al., 2009), there is notable Cre activity in the developing OS. This expression was evident during OS elongation in the dorsal side, which continues with the pRPE (E12.5, Fig. 1J, J’) and gradually expended ventrally as the optic fissure closes (E14.5, Fig. 1K, K’). By E14, after the closure of the optic fissure, we detected eGFP in the entire RPE, ciliary margin zone (CMZ), and OS derived astrocytes progenitors surrounding the RGCs axons (Fig. 1K). At P7, eGFP was detected in the RPE, pigmented cells of the ciliary body (CB), iris, ON astrocytes and some of the retinal astrocytes that likely migrate from the ONH into the GCL (Fig. 1L-L’). Thus, the *Dct-Cre* will delete the floxed genes in both pigmented and optic stalk lineages.

### Smarcc1 functionally replaces Smarcc2 in the pigmented and optic stalk lineages of the eye

The prominent expression of Smarcc2 in the RPE and ON astrocytes in the adult (Fig. 1F) inferred a key role for this subunit in their maturation and/or maintenance. We therefore examined the phenotype of *Smracc2^loxP/loxP^;Dct-Cre* (termed *Smarcc2-DCT-cKO*) adult mice eyes compared to control litter mates.

We did not observe, by histological analysis (H&E) of aged mice (P330) morphological differences in the OC derivatives; RPE, retina, optic nerve, CB and iris in the *Smarcc2-DCT-cKO* compared to the control litter mates (Fig. 2A, B). Although gross morphology and retinal thickness appeared preserved in the *Smarcc2-DCT-cKO* (Fig. 2C), more subtle alterations may result in reduced retinal function with age. However, electroretinography (ERG) analysis at P130, P260 and P380 revealed no significant differences in A- and B- wave amplitudes between the control and *Smarcc2-DCT-cKO* in both scotopic and photopic conditions (Fig. 2D, E, Fig. S2), indicating maintained retinal function despite Smarcc2 deletion in pigmented and optic stalk lineages.

**Fig. 2.**
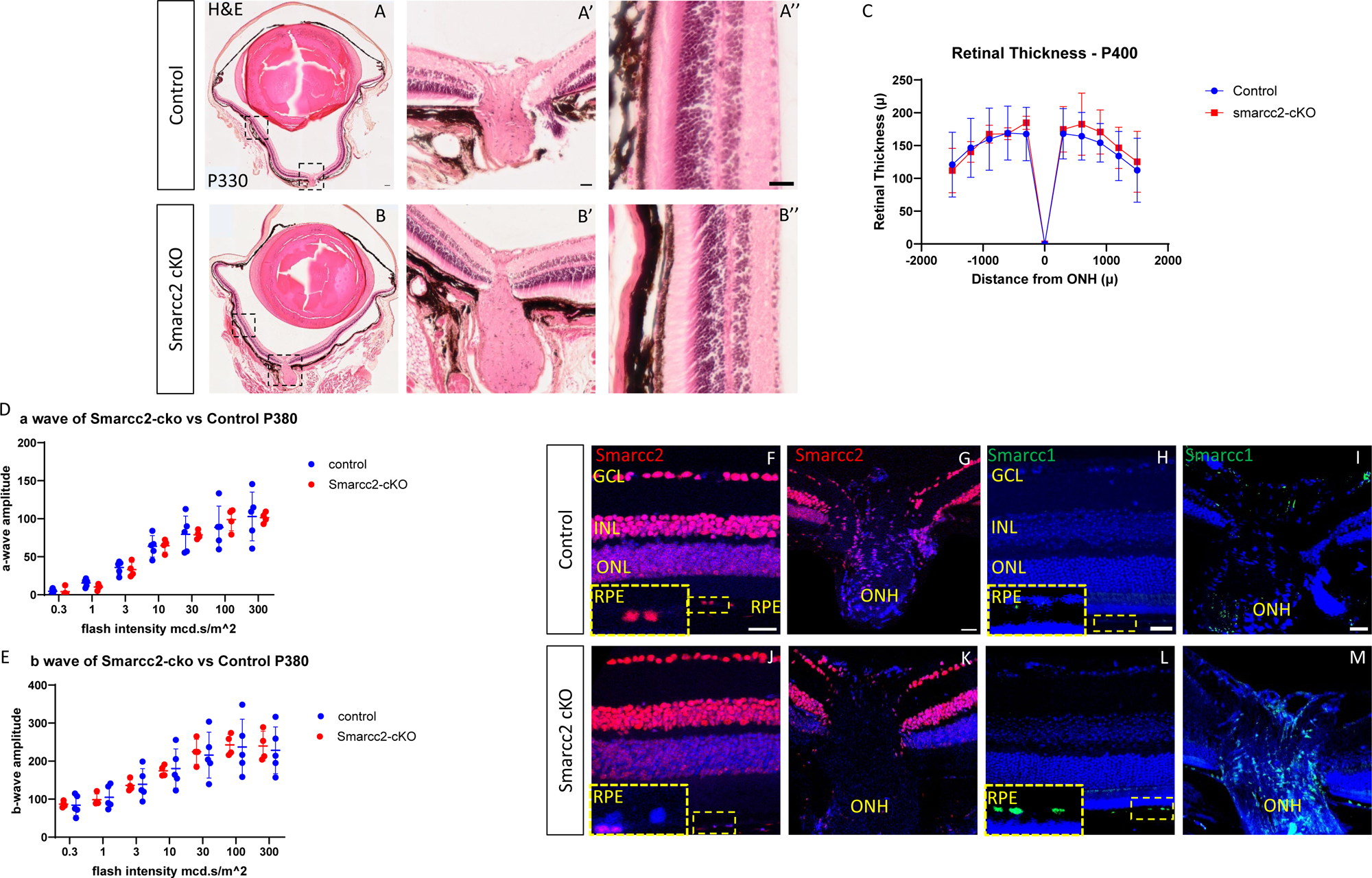
Smarcc1 functionally replaces Smarcc2 in the pigmented and OS lineages of the eye. Hematoxylin & Eosin (H&E) histology (A-B, larger magnifications of boxed region are in A’, A’’, B’, B’’) of the control (A-A’’) and *Smarcc2-cKO* (B-B’’) aged mice (P330). Retinal thickness measurements (C) of control and *Smarcc2-cKO* (n=3), five points for each side of the retina were measured with intervals of 300µ meters. Electroretinography (ERG) analyses for control and *Smarcc2-cKO* mice showing amplitude of a-wave (D) and b-wave (E) of scotopic test on aged mice (P380, n=4). Antibody labeling of control (F-I) and *Smarcc2-cKO* (J-M) aged mice (P380) for detection of Smarcc2 (F-G, J-K) and Smarcc1 (H-I, L-M). Scale bars A = 100µm, A’- I = 25µm.

The lack of detectable phenotype in the *Smarcc2-DCT-cKO* mice suggests functional compensation by Smarcc1. We examined the expression of both Smarcc1 and Smarcc2 comparing the control to *Smarcc2-DCT-cKO*, by antibody labeling. This analysis confirmed the selective expression of Smarcc2 but not Smarcc1 in the control RPE, retina and optic nerve head (Fig. 2F-I) and its selective deletion from the RPE and optic stalk in the *Smarcc2-DCT-cKO* eyes (Fig. 2J-K). The analysis further revealed an elevation of Smarcc1 protein in the RPE and ONH, where Smarcc2 was lost due to Dct-Cre activity (Fig. 2L-M). The observed elevation of Smarcc1 correlates with its capacity to functionally compensate for the loss of Smarcc2 in these cell types.

### Smarcc1 is required for optic nerve-head development but is redundant with Smarcc2 in the pigmented epithelium of the eye

We next examined the consequences of Smarcc1 deletion in the OS and pigmented epithelium lineages, in which Smarcc1 is co-expressed with Smarcc2 during embryogenesis (Fig. 1). In-vivo optical coherence tomography (OCT, A-D) of control (Fig. 3A, C) and *Smarcc1;loxP/loxP;DCT-Cre* (*Smarcc1-DCT-cKO*; Fig. 3B, D) demonstrated a severe malformation of the optic nerve head in Smarcc1-DCT-cKO old mice including atrophy of the adjacent retina and RPE as well as an abnormally excavated optic nerve head. These malformations were already evident by H&E analysis at P30 (Fig. 3 compare E, E’ control with F, F’’ of *Smarcc1-DCT-cKO),* whereas the pigmented lineages of RPE iris and ciliary body preserved their morphology and pigmentation (Fig. 3E’’, F’’). These results suggest that Smarcc2 can compensate for the loss of Smarcc1 in the development and morphology of the RPE, iris, and ciliary body but not in optic nerve head development. We further characterize the distribution of astrocytes and myelin in the ONH region of control (Fig. 3G) and *Smarcc1-DCT-cKO* (Fig. 3H) by antibody labeling for astrocyte marker GFAP and myelin basic protein (MBP). We observed a disorganized distribution of GFAP in the *Smarcc1-DCT-cKO*, while the separation between the myelinated and unmyelinated zones appears preserved in the *Smarcc1-DCT-cKO* mutants, as indicated by MBP staining, which identified myelin surrounding nerve fibers outside the unmyelinated zone in both control and mutant mice at P30 (Fig. 3G, H). These findings suggest that Smarcc1 is required for ONH morphology but not for the formation of an unmyelinated zone.

**Fig. 3.**
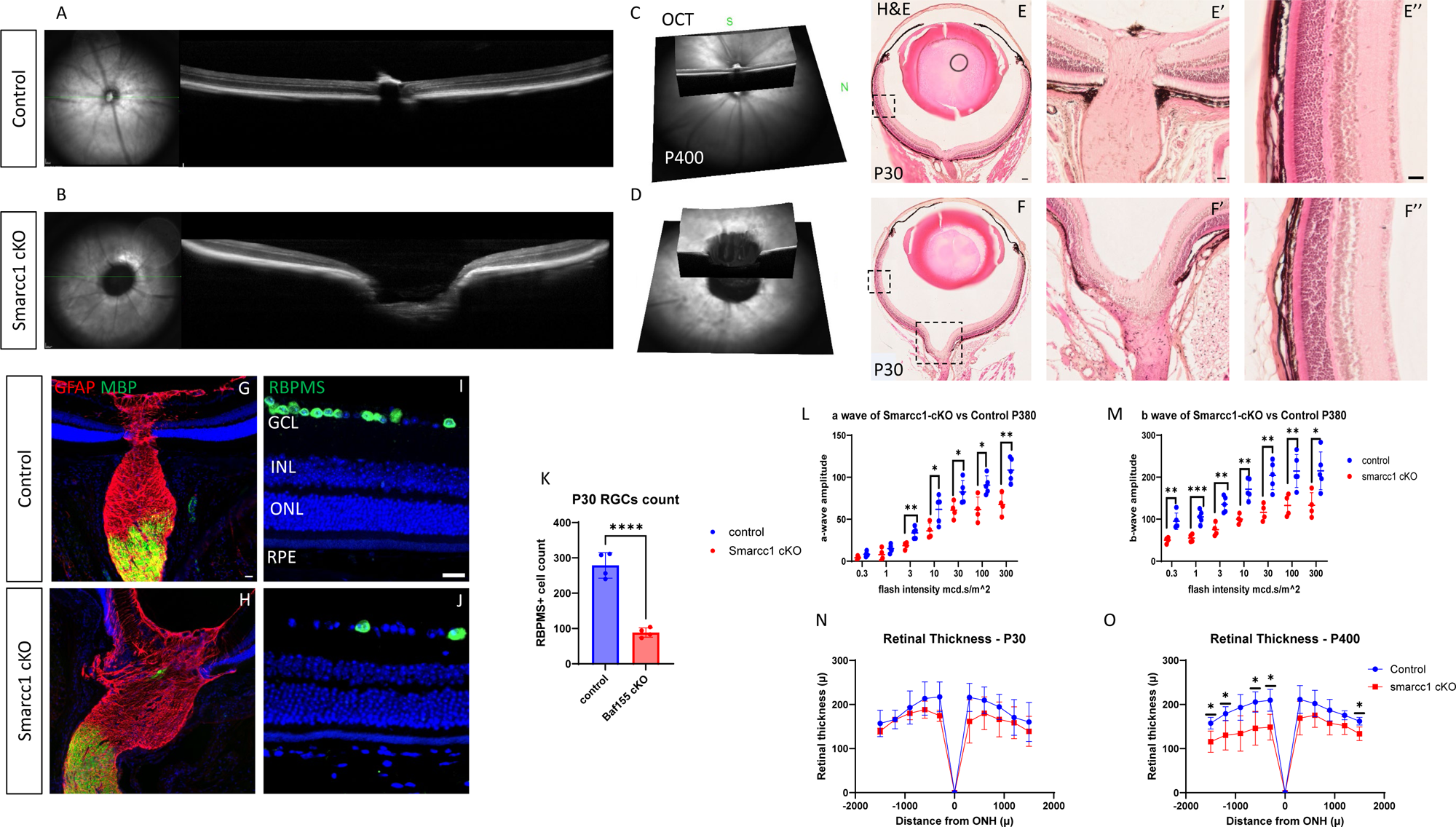
**Smarcc1 is required for optic nerve head development but functions redundantly with Smarcc2 in the pigmented epithelium of the eye**. Optical coherence tomography (OCT) imaging (A-D). Representative infrared and B-scan images of Control (A) and *Smarcc1-cKO* (B) retinas. OCT volumetric 3D renedring of optic nerve head area in control (C) and *Smarcc1-cKO*(D) mice. retinas, acquired at P400. Histology (H&E) of P30 Control (E, magnification in E’, E’’) and *Smarcc1-cKO* (F, magnification in F’, F’’) mice. Antibody labeling for Glial fibrillary acidic protein (GFAP), Myelin basic protein (MBP) (G,H), and RGCs marker RBPMS (I,J) in P30 control (G,I) and *Smarcc1-cKO* (H,J) mice. Quantification of number of RBPMS+ RGCs in P30 control and *Smarcc1-cKO* mice (K, n=4). Electroretinography (ERG) tests for Control and *Smarcc1-cKO* showing amplitude of a-wave (L) and b-wave (M) of scotopic test on aged mice (P380, n=4). Retinal thickness measurements of P30 (N) and P400 (O) Control and *Smarcc1-cKO* (n=3), five points for each side of the retina were measured with intervals of 300µm. Scale bars E = 100µm, E’- I = 25µm. T-test P-val *<0.05, **<0.01, ***<0.001, ****<0.0001.

The altered ONH morphology is likely to impact retinal ganglion cell (RGC) survival. Consistent with this, we observed a significant reduction in RBPMS-positive RGCs in *Smarcc1-DCT-cKO* compared with the control at P30 (Fig. 3I, J quantification in K). Although this reduction in RBPMS-positive GCL was not evident at P0, we did detect at this stage a significant increase in cleaved caspase 3 (CC3) mediated apoptosis in the *Smarcc1-DCT-cKO* GCL (Fig. S3E).

ERG analysis supports functional impairment, showing significant reductions in both A- and B-wave amplitudes in aged mice (P380) (Fig. 3L, M). In contrast, at earlier stages, ERG differences were not statistically significant, indicating a progressive decline in retinal function over time (Fig. S4). Furthermore, we detected a reduction in overall retinal thickness in the *Smarcc1-DCT-cKO* mice, which was more pronounced at P400 compared to P30, when the difference was not statistically significant (Fig. 3N, O).

These findings reveal a key role for Smarcc1 in ONH morphology. The reduction in retinal thickness and the progressive decline in retinal function likely primarily, but not exclusively, result from ONH abnormalities and surrounding retinal atrophy caused by Smarcc1 deficiency.

### The dorsal optic stalk resembles the pRPE progenitors of the optic cup

The phenotype in *Smarcc1-cKO* is primarily localized to the optic nerve head, which can be divided into two main regions. The first, termed the optic disc, is located at the site where axons exit the eye. The second, situated posterior to the optic disc (termed pONH), includes the glial lamina (lamina cribrosa in humans), a mesh of connective tissue originating from optic stalk astrocytes that supports the retinal axons as they exit the eye (Sun et al., 2009).

The morphogenesis of the optic fissure closure has been investigated primarily at the level of the optic cup, while the progression and regulation of this process at the level of the optic stalk remain mostly unresolved (Patel and Sowden, 2019). To further investigate the pattern of optic stalk maturation, we characterized, through antibody labeling, the expression of several early-expressed cell-type-specific TFs along the proximal-distal and dorsal-ventral axes.

Coronal sections presenting the dorsal and ventral optic stalk and cup of control embryos reveal that Pax6 is expressed throughout the early optic cup and also detected, albeit at lower levels, in the OS and the prospective optic disk (pOD) at E12 (Fig. 4A). At this stage, Otx2 and Mitf are confined to the dorsal layer of the OS that seems to continue with the pRPE of the optic cup whereas Pax2 expression is restricted to the ventral-proximal OS and the pOD, (Fig. 4B-D).

**Fig. 4.**
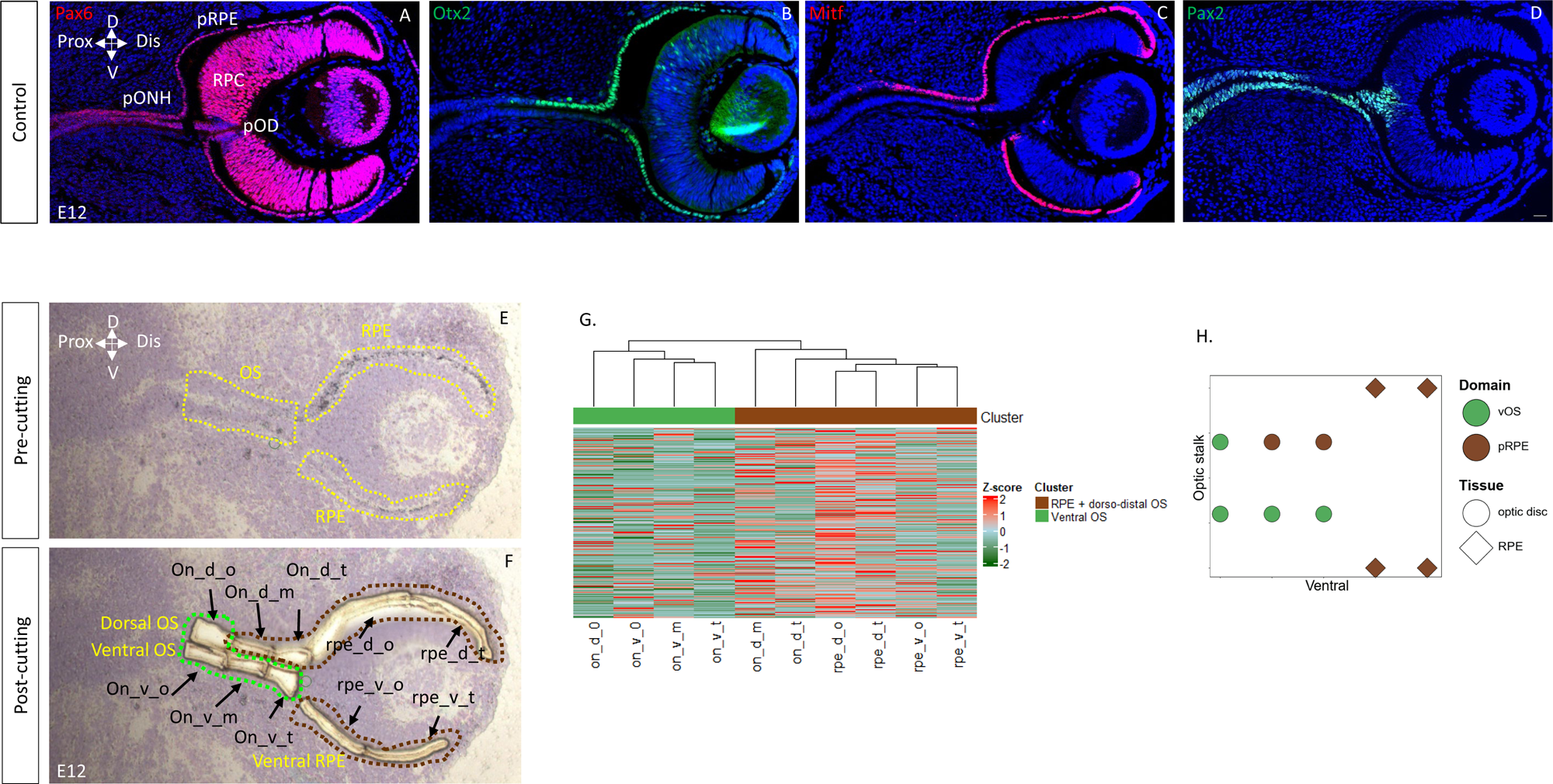
The dorsal optic stalk continues with the RPE progenitors of the optic cup. Antibody labeling for Pax6 (A), Pax2 (B), Otx2 (C), Mitf (D) in the E12 optic cup along dorsal-ventral axis. The GEO-Seq analysis on E12 cryosections was conducted on the indicated regions, isolated by laser capture microscope (LCM), along the dorsal ventral and proximal distal axes of the optic cup and optic stalk (before (E) and after (F) sectioning). Whole transcriptome normalized counts of each section (n=3-5) show two main groups as detected both in K-means clustering (K = 2) and with hierarchical clustering (G). Spatial plot illustrating unsupervised clustering of global gene expression, which divides the sample into two domains. The left side denotes the optic disc side, while the bottom represents the ventral side (H). The color of each domain is consistent with that in Panel G. Abbreviations: pOD - prospective optic disk, RPC – Retinal progenitor cells, pONH – Posterior optic disc, pRPE – Presumptive RPE. Scale bar = 25µm.

To characterize the global gene expression profile of the E12 OS and pRPE, we implemented geographical positional sequencing (Geo-Seq (Chen et al., 2017)) on E12 eyes (Materials and Methods). We sampled three separate sections from each of the dorsal and ventral regions of the E12 OS and two samples each from the dorsal and ventral E12 pRPE (Fig.4E, F). The spatial transcriptome of E12 OS and pRPE provides with ∼7M mapped reads, more than 60% mapping ratio, low mitochondrial gene percentage, and 3,000 genes per sample on average (Figure S5A, Table S1). We performed unsupervised K-means and hierarchical clustering on whole gene expression data to assess sample heterogeneity. Both methods consistently identified two distinct groups (Fig. 4G and scheme in H): The first includes the pRPE sections together with the two dorsal-distal OS samples (termed pRPE, labeled brown in G, H), while the second cluster included the ventral OS sections along with the most proximal dorsal OS section (termed vOS, labeled green in G, H).

These findings indicate that at E12, prior to optic fissure closure, the dorsal-distal OS shares transcriptional identity with the pRPE, suggesting a common developmental trajectory.

We next identified domain-specific signature genes (see methods) which are exclusively expressed in either the pRPE (3729 genes, p<0.01) or the vOS (202 genes, p<0.01). Domain-specific gene analysis revealed 3,729 pRPE-enriched genes and 202 vOS-enriched genes (p < 0.01; Fig. S5B, Table S2). GSEA of pRPE genes showed enrichment for forebrain development and light response pathways (Fig. S5C), consistent with the early stage of differentiation of these pRPE cells (Fig. S5C and Table S3).

### Smarcc1 is required for astrocyte progenitor specification from the dorsal optic stalk

Given the transcriptional differences between dorsal and ventral OS at E12, we next examined how Smarcc1 loss affects optic stalk differentiation and ONH development along the dorsal-ventral axis.

Using transverse sections at the pONH before and after optic fissure closure (E12, E14; Fig. 5), we characterized Smarcc1, Smarcc2 and the catalytic subunit Smarca4 expression in Smarcc1-DCT-cKO and control embryos. In controls, these subunits were detected in both pONH layers at E12 and E14 (Fig. S6A–F). In Smarcc1-DCT-cKO embryos, Smarcc1 was absent from the dorsal OS at E12 but retained in the ventral OS (Fig. S5G). By E14, Smarcc1 expression was lost throughout the OS (Fig. S5J). Notably, despite Smarcc1 loss, we did not detect at these stages change in the expression of Smarcc2 nor Smarca4 (Fig. S6H,I,K,L). These results define two critical stages for assessing Smarcc1 function: early dorsal OS cells at E12 and astrocyte precursors derived from them at E14.

**Fig. 5.**
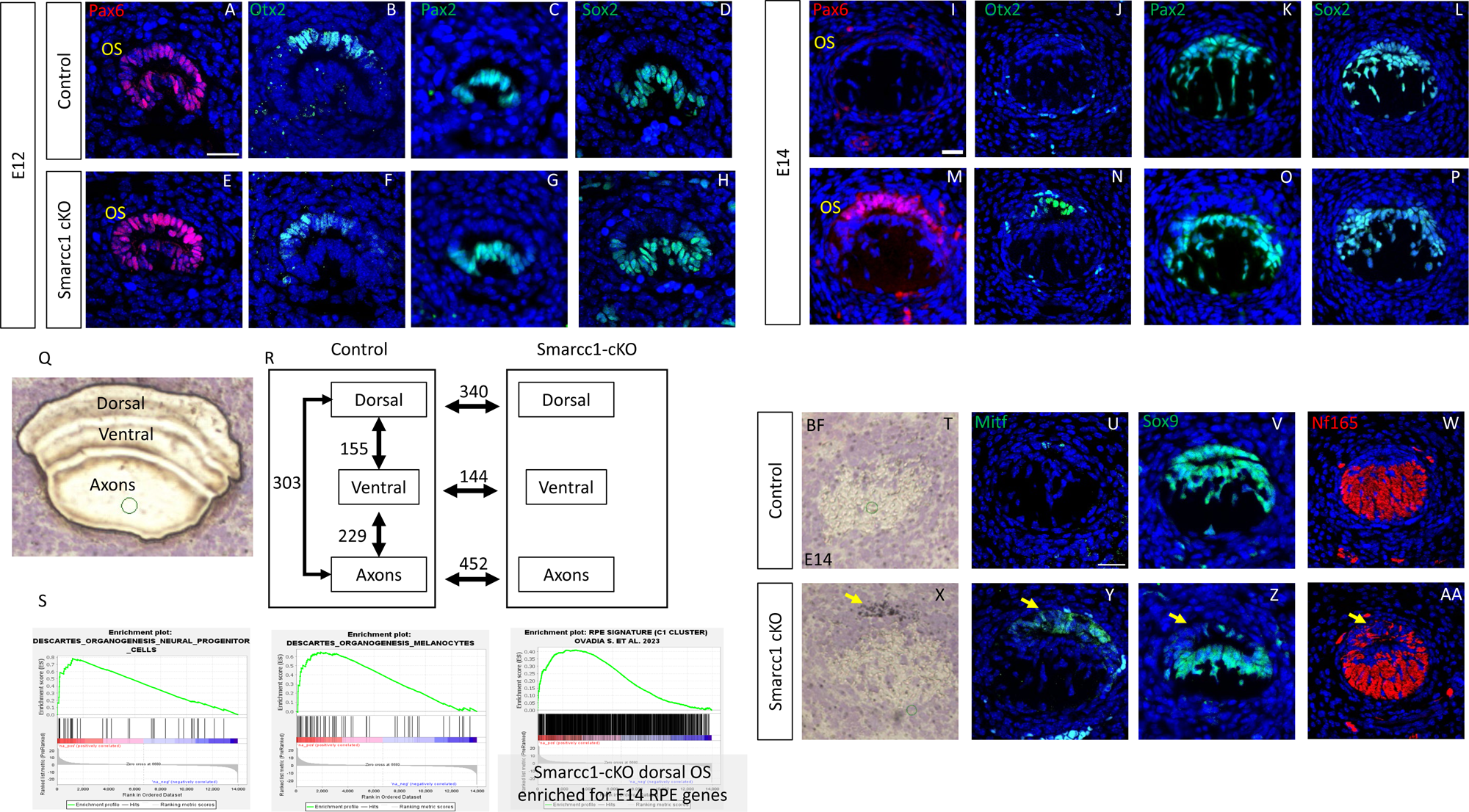
S**m**arcc1 **is required for dorsal optic stalk specification into astrocyte progenitors.** Cross sections of the OS of E12 (A-H) and E14 (I-P) mouse embryos stained with antibodies against Pax6 (A,E,I,M), Otx2 (B,F,J,N), Pax2 (C,G,K,O) and Sox2 (D,H,L,P). E14 OS section after LCM dissection of the three regions sampled (n = 3-5 samples per region). (Q). Scheme showing the number of DEGs between the compared regions and genotypes (R). GSEA shows that the Smarcc1-cKO dorsal region is enriched for neural progenitors, melanocytes, and E14 RPE signature genes (S). E14 OS cross sections of control (T-W) and Smarcc1-cKO (T’-W’) showing dorsal pigmentation (T-T’), and antibodies staining for Mitf (U-U’), NF165 (V-V’), Sox9 (W-W’). Scale bars = 25µm.

At E12.5, prior to optic fissure closure, Pax6 was detected in both dorsal and ventral OS, while Otx2 was restricted to the dorsal OS, consistent with its pRPE identity (Fig. 4A, B, Fig.5A, B). In contrast, the astrocyte progenitor transcription factors Pax2 and Sox2 were confined to the ventral OS (Fig. 4D, Fig. 5C, D). Despite the reduction of Smarcc1 in the dorsal OS of *Smarcc1-DCT-cKO* embryos, we did not detect a change in the expression levels of Pax6 or Otx2 at this stage (Fig. 5E-H), suggesting that at this stage the early dorsal OS identity is retained despite Smarcc1 loss.

By E14, following optic fissure closure, both Pax6 and Otx2 proteins were nearly undetectable in the OS (Fig. 5I-J, the gradual reduction of Otx2 in dorsal optic stalk is presented in Fig. S8). At this stage, astrocyte progenitors begin to delaminate from the dorsal OS and migrate into the forming optic nerve to ensheathe retinal axons (Fig. 5K-L). Accordingly, Pax2 and Sox2 expression expanded to both dorsal and ventral OS layers (Fig. 5K-L).

In contrast, *Smarcc1-DCT-cKO* embryos retained Pax6 and Otx2 expression in the dorsal OS at E14, while Pax2 and Sox2 were absent from this region (Fig. 5M-P). This suggests that the dorsal OS cells fail to initiate differentiation into astrocyte progenitors and instead maintain the early pRPE-like identity.

Together, these findings reveal a progressive differentiation pattern within the optic stalk and the role of Smarcc1 in both stages. At E12, dorsal and ventral OS cells exhibit distinct identities, with dorsal cells showing pRPE-like features. As development proceeds and the fissure closes, this distinction is lost, and all OS cells acquire astrocyte progenitor characteristics, marked by Sox2 and Pax2 expression. Smarcc1 is essential for this transition, particularly for the transition of dorsal OS cells from a pRPE-like state to astrocyte progenitors.

For global characterization of the differences between the dorsal and ventral regions of the OS during embryonic development and to reveal the transcriptional network governed by Smarcc1, we characterized the OS transcriptome at E14, by GEO-seq (Chen et al., 2017). This stage was chosen as the earliest stage a phenotype was detectable, and because the deletion occurs in all optic stalk cells by this stage.

Samples were collected by LCM from three regions of the E14 OS: dorsal, ventral, and RGCs axons (Fig. 5Q) from 4 control and 5 *Smarcc1-cKO* embryos, enabling a statistical evaluation of the variations between and within the samples.

From each sample we obtained, using Smart-Seq2, a mean depth of 30 million sequenced reads per sample. The raw reads were mapped onto the mouse reference genome (mm10) and an average of about 12,000 unique genes were identified per sample (Fig. S7A, Table S4).

Principal component analysis (PCA) presented high correlation between the samples isolated from the same regions (Fig. S8B). Notably the dorsal samples show separation in the PCA between the *Smarcc1-cKO* and the control, while in the ventral and axon regions of control and *Smarcc1-cKO* clustered together.

We determined the differentially expressed genes (DEGs) between the three regions, and between the control and *Smarcc1-DCT-cKO* samples of each region using DESeq2 [|log2(fold change)|>0.5, *P-*adj<0.05] ((Love et al., 2014), Fig. 5R, Table S5).

Gene Set Enrichment Analysis (GSEA) (Mootha et al., 2003; Subramanian et al., 2005) comparing gene expression between the dorsal Smarcc1-cKO and control dorsal regions reveals an enrichment for gene sets involved in melanocyte organogenesis and neural progenitor genes (Figure 5S; Table S6). Furthermore, GSEA of gene sets from E14 mouse pRPE, defined by Geo-seq (Ovadia et al., 2023) identified significant enrichment for pRPE signature genes in the E14 Smarcc1-cKO dorsal OS, but not in the ventral OS nor the axon-rich regions (Figure 5S, Table S6). This enrichment suggests that loss of Smarcc1 arrests dorsal OS cells at an early pRPE-like stage. Consistent with this, at E14 Smarcc1-cKO eyes—but not controls—displayed pigmented accumulation and Mitf expression, whereas Pax2 and Sox9, markers of astrocyte progenitors normally present in controls, were markedly reduced or undetectable (Fig. 5T–W; 5X–AA).

### Loss of Smarcc1 disrupts astrocyte progenitor proliferation dynamics in the optic stalk

The expansion of the dorsal optic stalk in Smarcc1-cKO pONH may result from altered cell proliferation, a process known to be regulated by SWI/SNF complexes in different lineages including the RPE (Braun et al., 2021; Elbaz-Hayoun et al., 2023; Ovadia et al., 2023). To investigate this, we assessed cell proliferation using a 5-ethynyl-2’-deoxyuridine (EdU) pulse-chase assay (3-hour pulse, (Salic and Mitchison, 2008)), quantifying EdU incorporation in both dorsal and ventral optic stalk cells (Fig. 6A, Fig. S9).

**Fig. 6.**
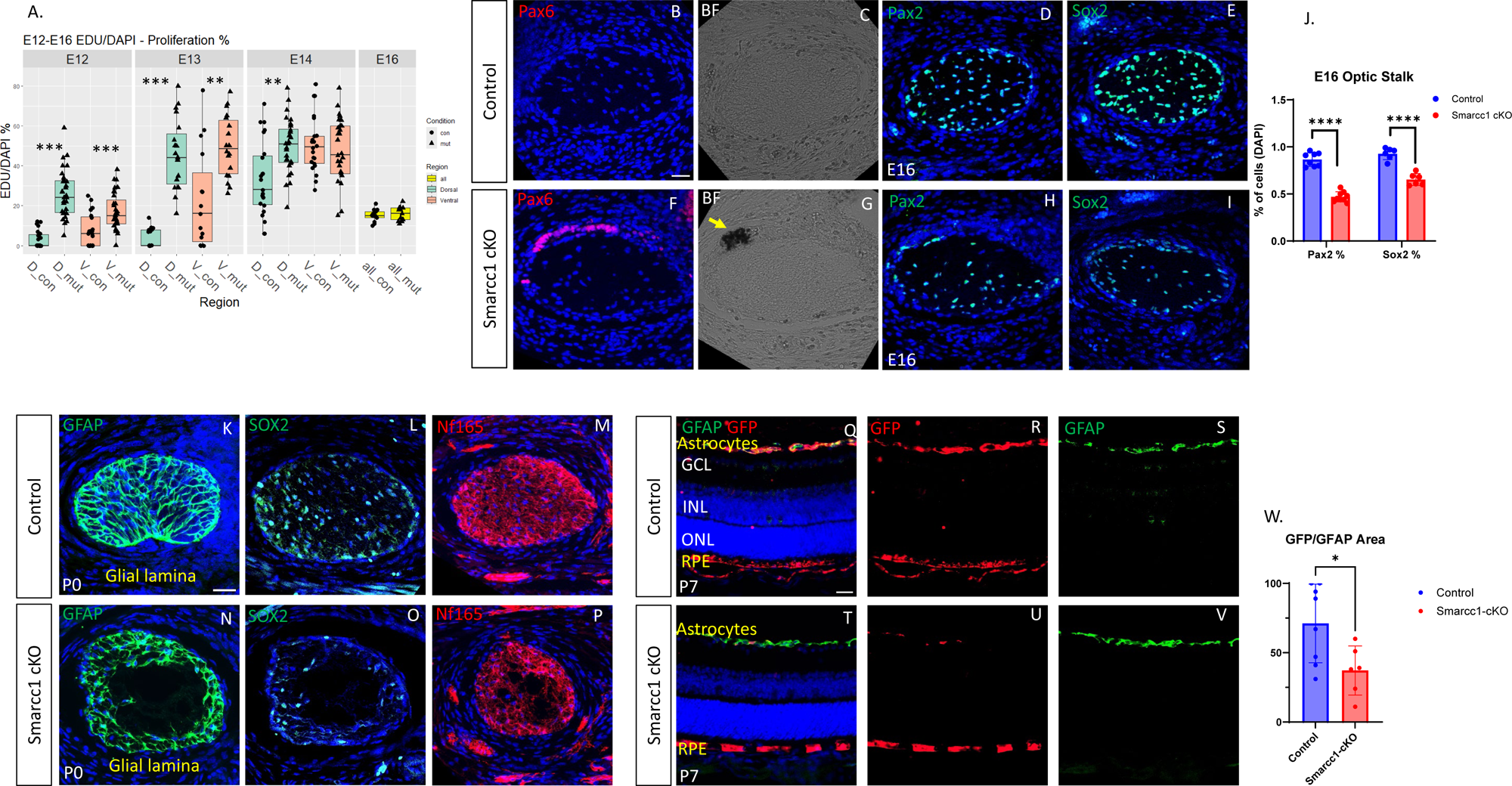
Smarcc1 is required for astrocyte progenitor proliferation dynamics, formation of the glia lamina and migration to the retina. OS cross-section quantification of EDU/DAPI in the dorsal and ventral regions of control (circles) and Smarcc1-cKO (triangles) following 3 hours pulse-chase at E12, E13, E14 and for E16 (without dorsal-ventral separation (n = 4-5) (A). E16 OS cross-sections of control (B-E) and Smarcc1-cKO (F-I) antibody staining for Pax2 (B,F), Sox2 (C,G), Pax6 (D,H) and bright-field (E,I). E16 OS Pax2 and Sox2 quantification (Target/DAPI) (n=4) (J). P0 OS cross-sections of control (K-M) and Smarcc1-cKO (N-P) antibody staining for GFAP (K,N), Sox2 (L,O), NF165 (M,P). P7 Retinal section of control (Q-S) and Smarcc1-cKO (T-V) labeled with GAFP and GFP, merged (Q,T) and separated channels (R,S,U,V). Quantification of the GFP positive area out of the GFAP area (GFP/GFAP) showing percentage of Smarcc1-cKO astrocytes in the retina (n=3) (W). Scale bars = 25µm. T-test P-val *<0.05, **<0.01, ***<0.001, ****<0.0001.

In control mice, the average EdU uptake in the dorsal optic stalk was approximately 5% of cells, at both E12 (3.04%, SD = 4%) and E13 (4.1%, SD = 4%), with a marked increase observed at E14 (33.2%, SD = 18%). In contrast, the ventral region, where the migrating astrocyte progenitors are located, showed a gradual increase in EdU-positive cells: 8% (SD = 9%) at E12, 23% (SD = 24%) at E13, and 53% (SD = 15%) at E14 (Fig. 6A).

In contrast in the Smarcc1-cKO mice, the dorsal optic stalk exhibited a significantly higher proportion of EdU-positive cells as early as E12 (26%, SD = 11%), increasing to 45% (SD = 17%) at E13 and 50% (SD = 13%) at E14 (Fig. 6A). Similarly, the ventral region showed accelerated proliferation, with 50% (SD = 18%) of cells EdU-positive already by E13—one day earlier than in controls (Fig. 6A).

These findings suggest that in control embryos, optic stalk cell proliferation is linked to the acquisition of astrocyte progenitor identity. Loss of Smarcc1 disrupts this coordination, leading to premature and excessive proliferation in both dorsal and ventral regions. Consequently, the average number of astrocyte progenitors in the Smarcc1-cKO pONH was significantly higher compared to the controls at E14 (Control: 24; SD = 5.7 cells, Smarcc1-cKO: 44 SD = 10.75 cells; Fig. S9).

By E16, the difference in cell proliferation between control and Smarcc1-cKO optic stalks is no longer observed. In both control and Smarcc1-cKO we detected 18% (SD = 2%), and 19% (SD = 3%) cells in S-phase (Fig. 6). This convergence suggests that the early proliferative surge in the Smarcc1-cKO optic stalk is transient and may be followed by a compensatory slowdown or normalization of cell-cycle activity.

### Smarcc1 is essential for ONH astrocyte differentiation and migration

At E16, the OS in the control embryos lacked Pax6 or pigment-expressing cells and astrocytes progenitors, marked by co-expression of Pax2 and Sox2, appeared evenly distributed around the RGCs axons (Fig. 6B-E). In contrast, the dorsal OS of the Smarcc1-cKO contained a small cluster of ectopic pigmented cells and Pax6 positive cells (Fig. 6F, G), likely remnants of dorsal OS cells that failed to acquire astrocyte progenitor identity due to Smarcc1 loss. Despite this, the total number of cells within the Smarcc1-cKO OS was comparable to the controls (82, SD = 9 in control and 89, SD = 18 in Smarcc1-cKO), but these cells were mostly confined to the OS periphery. Moreover, the proportions of Sox2 and Pax2 in these cells was significantly reduced (Pax2 - Control -86%, SD = 3%. Smarcc1-cKO – 46%, SD = 3%. Sox2 - Control -92%, SD = 3%. Smarcc1-cKO – 65%, SD = 3% Fig. 6H-J). Thus, Smarcc1 loss leads to diminished expression of astrocyte progenitors TFs and impaired migration into the developing ON, indicating failure in astrocyte progenitor maturation.

ONH astrocytes play a critical role in maintaining structural integrity for RGC axons exiting the eye by forming the glial lamina—a dense astrocytic meshwork that ensheathes RGC bundles in the unmyelinated prelaminar pONH. In controls at P0, this meshwork expressed Sox2 and GFAP and surrounded retinal axons labeled by Nf165 (Fig. 6K–M). In Smarcc1-cKO mice, however, astrocytes failed to migrate into the pONH, remained at the periphery, and exhibited reduced Sox2 expression (Fig. 6N, O). Correspondingly, RGC axons were depleted in astrocyte-deficient regions (Fig. 6P), consistent with RGC death observed at this stage (Fig. S3). These findings highlight Smarcc1’s essential role in astrocyte differentiation, glial lamina formation, ONH integrity, and postnatal RGC survival. To further assess Smarcc1’s role in ONH astrocyte migration into the retina, we traced the astrocytes in which DCT-Cre was active by utilizing the Z/EG reporter (Novak et al., 2000). This analysis revealed that DCT-Cre transgene was active in some, but not all, Pax2-positive optic disc (Fig. S10). In the P7 control (DCT-cre;Z/EG) retinas, eGFP was detected in approximately 70% (SD = 26%) of retinal astrocytes in the RGC layer indicating their origin in the ONH (Fig. 6Q-S).

In Smarcc1-cKO mice, this proportion was significantly reduced to ∼37% (SD = 16%) (Fig. 6T–W), suggesting impaired astrocyte migration from the ONH into the retina.

Together, these results demonstrate that Smarcc1 is required for the proper development, organization, and migratory capacity of ONH astrocytes. Its loss disrupts glial lamina formation and astrocyte positioning, compromising ONH integrity and leading to RGC degeneration.

## Discussion

### Dorsal-ventral patterning and temporal regulation of astrocyte differentiation in the optic stalk

Our findings reveal a previously underappreciated dorsal-ventral axis pattern of astrocyte cell-fate acquisition within the optic stalk, which complements the well-established proximal-distal axis that delineates the optic stalk, neural retina, and RPE lineages. Traditionally, proximal-distal patterning of the optic vesicle has been attributed to boundary formation mediated by cross-inhibition among region-specific TFs, regulated by signaling pathways such as Sonic Hedgehog (Shh) and the Tgfb (reviewed in (Bharti et al., 2006; Fuhrmann et al., 2014; Moreno-Marmol et al., 2018; Tao and Zhang, 2014)). By analyzing this process before (E12) and after (E14) optic fissure closure using antibody labeling and spatial transcriptomics, we found that the dorsal and ventral regions of the optic stalk differ in both molecular identity and timing of cell fate specification. The dorsal stalk, adjacent to the outer layer of the optic cup, expresses early RPE-associated TFs—Pax6, Otx2, Mitf—suggesting RPE-like properties, while the ventral stalk, which is continues with the neuroretina progenitors in the inner optic cup, initiates astrocyte differentiation earlier, marked by expression of Pax2 and Sox2 early markers for type 1 astrocyte progenitors of the ON. These markers appear in the ventral stalk around embryonic day 12 (E12) and extend dorsally by E14, indicating a wave of differentiation that mirrors the spatial maturation of the optic stalk (Fig. 7 A-B).

**Fig. 7.**
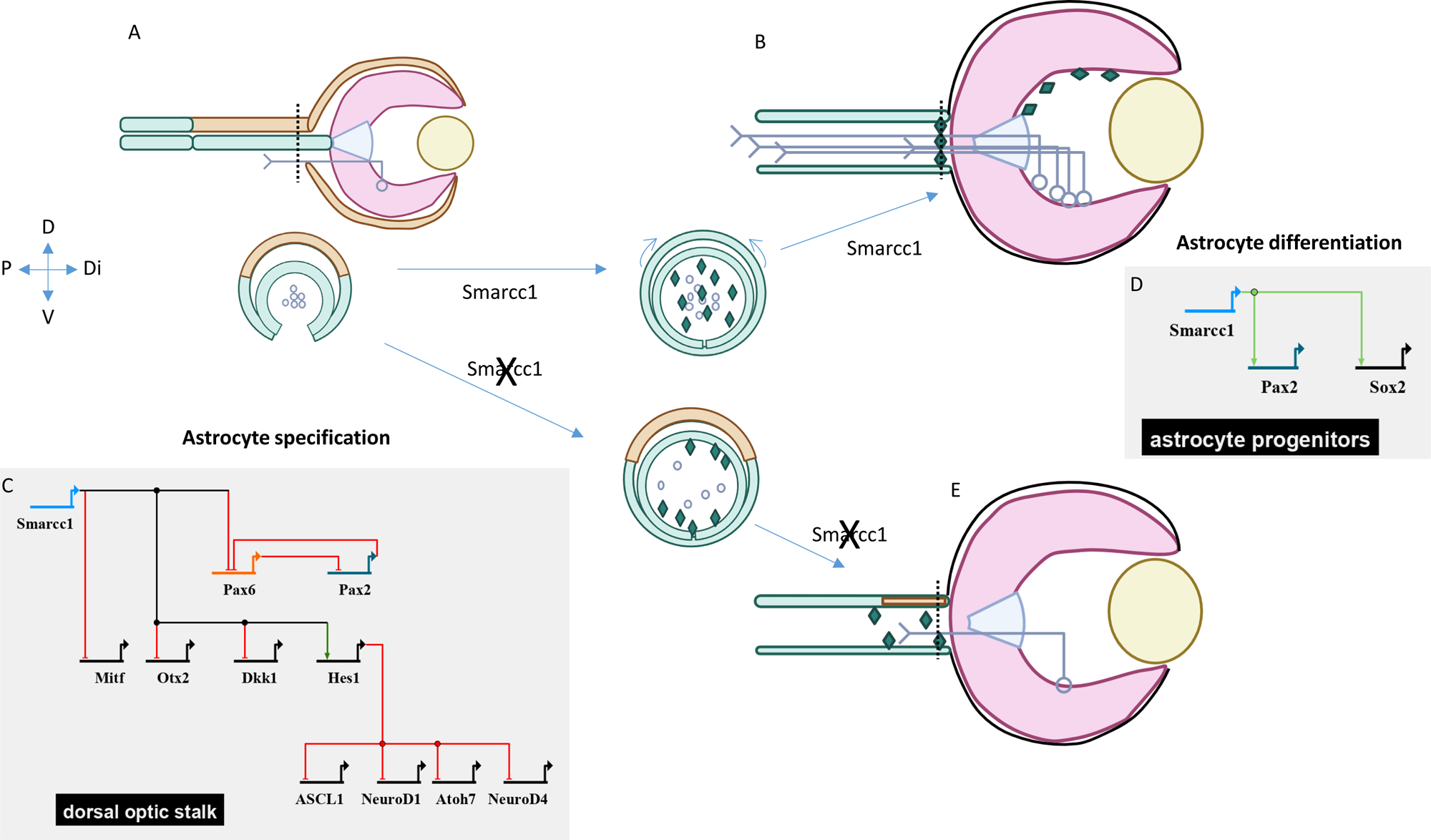
S**p**atiotemporal **specification and differentiation of optic nerve astrocytes from optic stalk neuroectoderm and their dependence on Smarcc1.** (A) Early stage (E12, mouse): dorsal optic stalk retains pigmented progenitor identity. (B) Later stage (E14, mouse) following completion of the ventral-to-dorsal wave of astrocyte specification (arrows) marked by Pax2 and Sox2, followed by delamination and migration of astrocyte progenitors (diamonds) into the optic nerve to form the glial lamina. The cross section along the optic stalk in each stage is presented below. Smarcc1 activity is required for astrocyte specification from dorsal optic stalk through downregulation of pigmented epithelium determinants (Pax6, Otx2, Mitf) and expression of astrocyte markers (Pax2, Sox2, Sox9). Loss of Smarcc1 during astrocyte progenitors differentiation disrupts glial lamina formation and optic nerve head morphology, leading to ganglion cell death. The GRNs for each stage is presented based on spatial transcriptomic analysis (BioTapestry (Longabaugh et al., 2005) Illustrations created in BioRender.

Notably the ventral to dorsal wave of astrocyte specification is tightly coupled with two key developmental events: optic fissure closure, which is completed by E12.5, and axonal extension from retinal ganglion cells toward the brain. The dorsal stalk is the last to differentiate into astrocyte progenitors, suggesting a spatially coordinated maturation process (Chan et al., 2020). In this regard, the inner layer of the optic stalk, which interfaces with extending axons, appears to proliferate more and delaminate prior to the outer layer, potentially regulating both the timing and quantity of astrocyte production. This layered maturation may reflect a mechanism to ensure that astrocyte differentiation is synchronized with axonal pathfinding and optic nerve formation (Huxlin et al., 1992; Tao and Zhang, 2014). Such coordination is critical, as astrocytes not only support axonal growth but also contribute to the formation of the gliovascular network and optic nerve architecture.

The transient RPE-like identity of the dorsal stalk aligns with previous observations of ectopic pigmentation following genetic perturbations that disrupt fate determination of RPE and retinal lineages. For instance, Pax2 knockout or mutations in Vax1/2 result in expansion of both retina and RPE territories, abolishing optic stalk formation (Hallonet et al., 1999; Mui et al., 2005; Torres et al., 1996). Another example involved the misexpression of Pax6 in the optic stalk, using Pax2 enhancer. This perturbation was expected to result in replacement of stalk with ectopic retina and RPE, yet the outcome was restricted to pigment accumulation along the dorsal stalk (Schwarz et al., 2000). These findings underscore the differential molecular properties of the dorsal and ventral optic stalk and their distinct responses to genetic perturbations.

Our data further indicates that the posterior optic nerve head region is delayed in acquiring astrocytic identity, and instead retains RPE-like transcriptional features after the adjacent optic stalk regions have committed to the astrocyte lineage. This, which is based the pattern of expression of Pax6, Mitf, Otx2 (Figs 4, S8, Fig. 7 A-B), suggests that the posterior ONH may represent a transitional or regulatory zone, where cell fate decisions are temporally postponed, possibly to coordinate with late-stage events such as optic cup morphogenesis, vascular invasion, and glial lamina formation as well as postnatal growth at the posterior optic nerve head (Bernstein et al., 2020). Taken together, the optic nerve head exhibits a delayed onset of astrocyte marker expression compared to the rest of the optic stalk. This region retains RPE-like transcriptional signatures late into development, suggesting a unique temporal regulation of fate specification that may reflect its specialized role in optic nerve formation and function.

### Functional redundancy and divergence of Smarcc1 and Smarcc2 in eye development

Within this spatial framework, our data highlight distinct roles for Smarcc1 and Smarcc2, the core components of the SWI/SNF complex. In Smarcc2-deficient models, we observe upregulation of Smarcc1, with no apparent disruption in the morphology or function of the RPE, optic stalk, iris, or ciliary body. This suggests that Smarcc1 is sufficient to compensate for Smarcc2 loss, consistent with prior studies indicating functional redundancy between these subunits in maintaining BAF complex functions (Fig. 2, (Narayanan et al., 2018)). The expression of Smarcc2 in the pigmented lineages of the eye is nevertheless important for compensating for Smarcc1 loss in the RPE, iris and ciliary body lineages as these structures seem morphologically intact in the Smarcc1-cKO (Fig. 3). Thus, pigmented lineages derived from the outer optic cup seem to require either Smarcc1 or Smarcc2, which together ensure robust differentiation of these tissues.

### Smarcc1 is obligatory for OS patterning and for OS astrocytes functions

Nevertheless, the loss of Smarcc1 does expose its non-redundant role in optic nerve head development, a pivotal structure for retinal function. In the Smarcc1 conditional inactivation we detected optic nerve head dysgenesis resulting in capping of the optic nerve head. The alterations in the optic nerve head seem to result in non-cell-autonomous effect on the retina, as it was accompanied by early loss of ganglion cells and reduced retinal activity based on ERG analysis in aged mice.

Detailed characterization of DCT-Cre activity onset shows that Smarcc1 deletion begins at E12 in the dorsal optic stalk, prior to astrocyte specification, and expands ventrally by E14, affecting cells already committed to the astrocyte lineage. Phenotypic analysis indicates that Smarcc1 is essential at both stages. First, it is required for the specification of dorsal optic stalk cells into astrocytes. In its absence, these cells fail to initiate astrocyte differentiation and instead retain progenitor-like, RPE-like characteristics, marked by elevated proliferation and expression of Mitf, Otx2, and Pax6. Concurrently, these cells show reduced expression of Hes1 and upregulation of proneural genes further suggests a failure to suppress nuroepithelial identity and promote glial fate, resulting in the co-expression of competing RPE and neural precursor genes (Fig. 7). Although dorsal cells initially expand due to increased proliferation, this is not sustained, and the pigmented cell patch observed at P0 is small and does not integrate into the ONH structure.

In contrast, ventral optic stalk cells in the Smarcc1-cKO model do acquire astrocyte progenitor identity before Smarcc1 deletion occurs and thus providing opportunity to study Smarcc1 loss in course of astrocyte maturation. By E16 the distribution of OS cells originating from the ventral side seems altered, with more cells at the periphery of the OS and less delaminated between the RGCs axons. By P0, these astrocyte progenitors populate mostly the periphery of the optic nerve and at this stage we detect regions which are devoid of axons, based on Nf165 distribution (Fig. 6). Furthermore, the Smarcc1 astrocytes seem to fail to migrate into the nerve fiber layer of the retina. These results show that Smarcc1 is not only obligatory for OS cells to gain astrocyte identity, but also for normal astrocyte differentiation in respect of migration into the optic nerve and to the retina. The detected reduction in Sox2 already at E16, may explain some of these phenotypes as in the Sox2 is required for astrocyte differentiation in the retina and it functions in a unique subset of optic nerve head astrocytes for postnatal optic nerve growth (Bernstein et al., 2020; Kautzman et al., 2018; Wang et al., 2022). The transcriptomic analysis of the Smarcc1-cKO further revealed reduction in expression of ECM modulators such as Mmp14 and Adam15, which were shown to control cell migration (Garmon et al., 2018; Martin et al., 2002). Collectively, Smarcc1 seem to be upstream of multiple pathways contributing to astrocytes identity and migration properties.

### Smarcc1 loss in retinal astrocytes is compensated by normal cells

The formation of the astrocyte template and the regulation of astrocyte numbers in the retinal nerve fiber layer (RNFL) is a tightly controlled developmental process, critical for ensuring proper retinal vasculature patterning and integrity (Paisley and Kay, 2021). During development, an excess of astrocytes is produced, far greater than the number present in the mature retina. Notably, recent studies show that microglia play an active role in this regulation by selectively removing surplus astrocytes through non-apoptotic mechanisms, a process essential for normal reginal vascular development (Punal et al., 2019). In our study, we observed that Smarcc1 loss reduces the number of mutant astrocytes, but total astrocyte counts in the retina remain unchanged. This suggests that genotypically normal astrocytes—likely sourced from the retinal disc—can compensate for the loss of a mutant subpopulation. Such compensation demonstrates cellular plasticity and provides direct evidence for robust homeostatic mechanisms that preserve the astrocyte template despite perturbations. These findings underscore the broader principle of glial compensation, which may have relevance not only during development but also in the context of retinal disease, repair, or regeneration.

## Materials and methods

### Animals

Mice were kept at the Tel Aviv University Animal House Facility. Animal use was approved by the Tel Aviv University Animal Care and Use Committee (Approval Number 28-02-2021). The analysis was conducted on Baf155^loxP/loxP^, Dct-Cre;Z/EG or Baf170^loxP/loxP^;Dct-Cre;Z/EG and control littermates that do not carry the DCT-Cre (the genotypes of the GEO-seq samples are listed in Table S1). The Baf155^loxP/+^, Baf170^loxP/+^, Dct-Cre and Z/EG mouse lines (Novak et al., 2000) (Choi et al., 2012) (Davis et al., 2009) (Tuoc et al., 2013a) were maintained on a C57Bl6/J genetic background. The primers used for genotyping are listed in Table S7.

### Electroretinography

ERG recordings were performed in a dark room following overnight dark adaptation. Recordings were conducted using the portable HMsERG unit (OcuScience, Henderson, NV). Pupils were dilated using 0.5% tropicamide (Mydramide, Dr. Fischer), and 20 min later animals were sedated by a subcutaneous injection of ketamine and xylazine and placed inside a faraday cage to minimize noise interference. An active mouse contact lens electrode (OcuScience, Henderson, NV) was placed on the cornea, coupled with methylcellulose. Two subcutaneous needles were placed at the lateral canthus and base of the tail and served as reference and ground electrodes, respectively. Impedance was kept below 5 kΩ. Scotopic (dark adapted) function was recorded in response to 0.3, 1, 3, 10, 30, 100 and 300 mcd*s/m^2^ flashes in a randomly-determined eye. Following light adaptation (10 min, 30 cd/m^2^) photopic function was recorded bilaterally in response to 10, 30, 100, 300, 1000, 3000, 10000 and 25000 mcd*s/m^2^ flashes.

### Histology, immunofluorescence and EdU analyses

The embryonic heads were fixed with 4% (v/v) paraformaldehyde for 3 h and embedded in paraffin. Immunofluorescence analyses and hematoxylin and eosin staining were performed on 10-μm paraffin sections as described (Ashery-Padan et al., 2000). Briefly, sections were immersed in PBS and boiled twice in Unmasking Solution (VECTOR, VE-H-3300). The sections were blocked for 2 h in PBSTG (0.2% Tween 20, 0.2% gelatin in PBS), incubated overnight with primary antibodies, washed with PBSTG, incubated for 2 h at room temperature with the secondary antibodies, washed with PBSTG, and mounted in fluorescent mounting medium (OriGene E19-18). The source of antibodies and dilutions are listed in Table S8.

EdU pulse-chase experiments were done by intraperitoneal injection of EdU 50µg/kg body weight (Lumiprobe 0540) to pregnant mice three hours before sacrifice. EdU staining was done by click-chemistry method. The slides were incubated for 35 min before the secondary antibody in solution containing CuSO4, L-ascorbic acid (A4544), and Sulfo-cyanine-azide (Lumiprobe 2330).

For quantification the percentage of EdU from total DAPI positive cells was determined for the dorsal and ventral regions of interest (Fig. S9). The statistical analyses were performed using GraphPad Prism multiple T-test as detailed in the figure legends.

Retinal thickness was measured from the H&E staining of each cKO (n=3) (Smarcc1 or Smarcc2) and its corresponding control (n=3). Ten measurement points were evenly spread at 300 micron steps, five for each side of the retina, beginning from the ONH. The measurement included the entire retina, from the RPE to the RNFL. The imaging was conducted with an Olympus BX61 fluorescence microscope or an Andor BC43 confocal microscope.

### Optical coherence tomography

Optical coherence tomography (OCT) was performed to obtain high-resolution, cross-sectional images of the mouse retina. Mice were anesthetized with an intraperitoneal injection of ketamine (85 mg/kg, Vetoquinol, Lure, France) and the relaxing agent xylazine (15 mg/kg, Eurovet Animal Health, Bladel, the Netherlands), and pupils were dilated with Cicloplegicedol (cyclopentolate hydrochloride 1%, Laboratorios Edol, Linda-a-Velha, Portugal) prior to imaging. Retinal scans were acquired using a Heidelberg Spectralis OCT system (Heidelberg, Germany) equipped with a mouse imaging adapter. For each eye, an en-face infrared image of the posterior pole of the eye as well as a series of B-scans covering the central retina were collected in high resolution (HR) mode, providing an axial resolution of up to 3 μm and a lateral (transverse) resolution of approximately 5.7 μm.. Volumetric datasets were reconstructed from the acquired scans using the manufacturer’s software and further processed for three-dimensional rendering of the retinal layers.

### Geo-Seq

The spatial transcriptome of the developing RPE and OS was obtained according to the Geo-seq (Chen et al., 2017) protocol with minor modifications. Briefly, E12 and E14 mouse embryonic heads were embedded in OCT (Leica Microsystems, 020108926) and cryosectioned at a thickness of 12 μm. Sections were transferred onto LCM PEN membrane slides, immediately fixed in ethanol, and stained with 1% (v/v) Cresyl Violet acetate in 75% ethanol solution (Sigma-Aldrich).

From the selected sections, using LCM, we isolated four RPE sections and six OS sections of the E12 embryos (four controls and six Smarcc1-cKO), or three sections of the transverse OS from the E14 embryos (four controls and five Smarcc1-cKO). The samples were collected by laser microdissection, and total RNA pellets were dissolved in a lysis solution, followed by reverse transcription using SuperScript II reverse transcriptase (Invitrogen), and whole transcription amplification with KAPA HiFi HotStart ReadyMix (2×; KAPA Biosystems).

PCR products from the LCM samples were used for RNA-seq library construction. Briefly, PCR products were purified using 0.8:1 ratio AMPure XP beads (Agencourt), quantified with Qubit dsDNA HS Assay Kit (Thermo Fisher Scientific) on an Agilent Tapestation 4200, and a cDNA library was constructed using the Nextera XT Library Prep Kit (Illumina). The E12 samples were sequenced with Novaseq 6000 and the E14 were sequenced on an Illumina Nextseq 2000 instrument both using the 150-bp single-end reads setting.

### Pre-processing of Geo-seq data

The E12 sequencing quality of all raw sequencing data were evaluated by FASTQC and were separately mapped to the mouse GRCm38 genome assemblies using HISAT2 (Pertea et al., 2016) using default settings. Mapping ratio was calculated based on the number of mapped reads and total reads for each sample. All mapped reads were processed by Stringtie (Pertea et al., 2016) to quantify gene expression levels (measured by TPM, Transcripts Per Kilobase per Million mapped reads) using default parameters.

The E14 RNA-seq data were preprocessed using the Seq2Science pipeline (van der Sande et al., 2023) with the default settings. Briefly, we aligned the reads to the mm10 mouse reference genome using STAR (Dobin et al., 2013). Gene counting was done using HTSeq (Anders et al., 2015), and transcripts per million (TPM) normalization was done using Salmon (Patro et al., 2017).

### Identification of domain-specific expressed gene

To quantify the domain specificity of a gene, an entropy-based strategy was modified from our previous work (Cui et al., 2022). The Jensen–Shannon divergence (JSD) algorithm was used to identify spatial domain-specific genes by philentropy package (version 0.5.0) (https://doi.org/10.21105/joss.00765). For distribution of each gene P1 and predefined pattern P2, the spatial domain- specificity score (SSS) between P1 and P2 was defined by converting JSD to a similarity score:

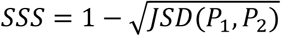

To evaluate the significance of gene specificity in each spatial domain, the spatial domain-specificity score was permuted across all samples 1,000 times, then calculated SSS, and finally a permutation p value was calculated as the number of times that *SSS*permutation > *SSS*true divided by 1,000. Only the significant genes (P < 0.01), which were specifically enriched in spatial domains, were kept for downstream analysis. We used GSVA (Hanzelmann et al., 2013), a unsupervised method, to estimate variation of domain-specific genes enrichment through the samples of a expression data set.

### DEGs and clustering analysis

Differentially expressed gene analysis was performed using DESeq2 (Pertea et al., 2016) (Love et al., 2014). The count matrix was prefiltered to include only genes with more than 10 counts in at least three samples and then subjected to library size normalization and variance stabilizing transformation (VST). The normalized transformed data were then subjected to dimensionality reduction using Principal Component Analysis (PCA), and to correlation analysis following hierarchical clustering using the stats and pheatmap package in R to assess sample quality and variance.

To infer all spatial and region-level DEGs, the following statistical tests were performed: (1) each region was compared with all other regions (Dorsal vs Ventral, Dorsal vs Axons, Ventral vs Axons); (2) for each of these regions, a comparison was performed between the control vs the Smarcc1-DCT-cKO. Genes with adjusted *P*-value lower than 0.05 and log2 (fold change) higher/lower than 0.58/−0.58 were considered DEGs. E12 gene expression clustering was performed on the normalized gene counts with both hierarchical clustering and K-means clustering separately, grouping the samples into the same two groups.

### GSEA of region comparison

The genes were ordered based on their log2(fold change) from the analysis, and were subjected to GSEA software (Subramanian et al., 2005) (Mootha et al., 2003). A minimum gene set size of 15 and maximum gene set size of 5000 genes were used, with a *P*-value cutoff of 0.05 and default false discovery rate (FDR).

For the GESA comparison between the E14 OS samples and the E14 RPE control and FcKO samples from our previous paper we used the K-means clustered DEGs representing the control E14 RPE from the RPE vs mesenchyme comparison, and the DEGs clusters representing control RPE or FcKO RPE from the comparison of those to conditions (Ovadia et al., 2023).

## Legends to Supplementary Figures and Tables

**Fig. S1.**
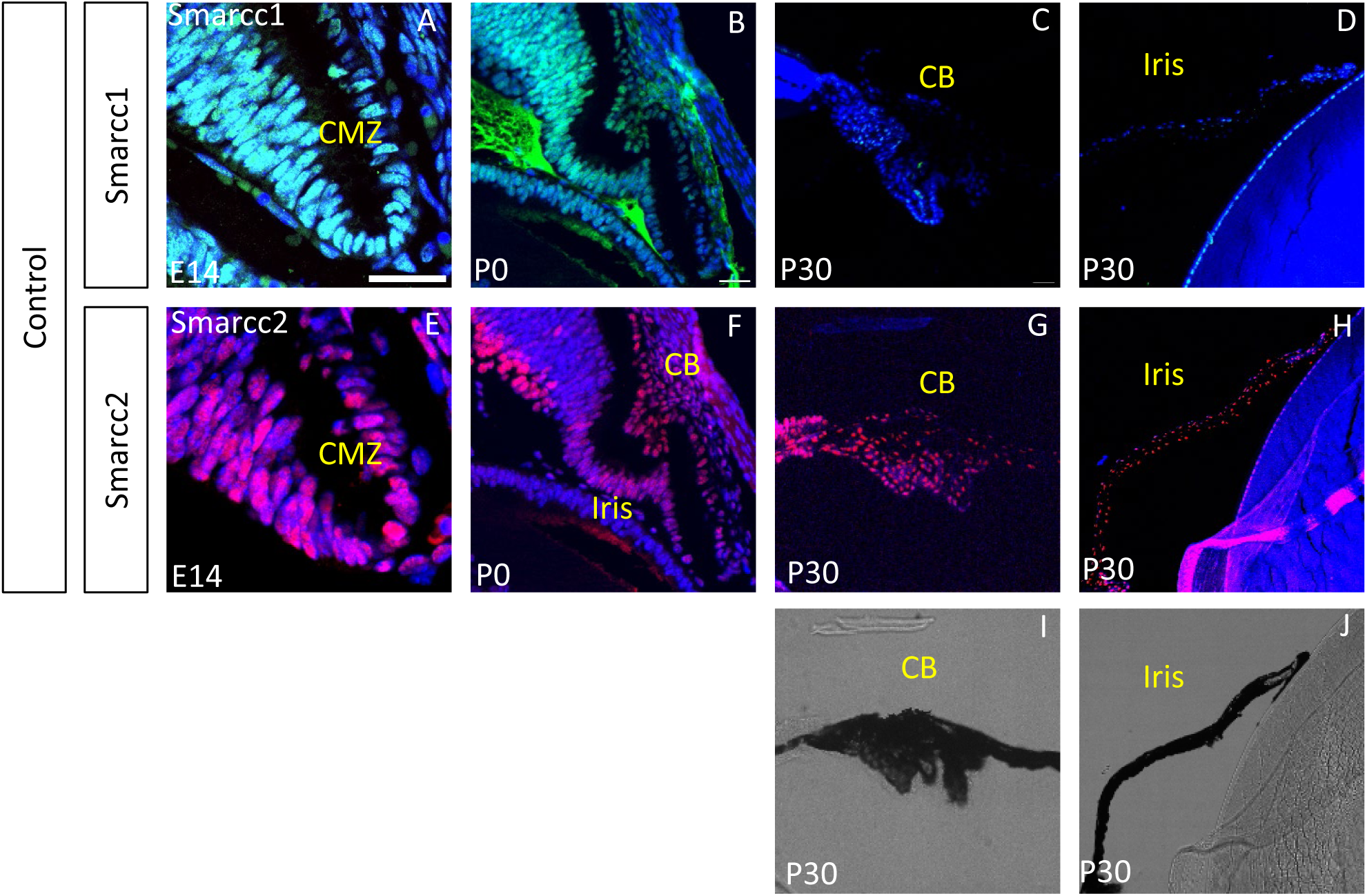
The stage-specific expression of Smarcc1 and Smarcc2 in the ciliary margin lineages. Antibody labeling for Smarcc1 (A-D), Smarcc2 (E-H) of sections of the ciliary margin at E14 (A, E), P0 (B, F) and iris and ciliary body at P30 (C, D, G, H), including the respective bright field view of the P30 iris (I) and ciliary body at P30 (J). Abbreviations: CMZ-Ciliary Margin Zone, CB-Ciliary Body. Scale bars = 25µm.

**Fig. S2.**
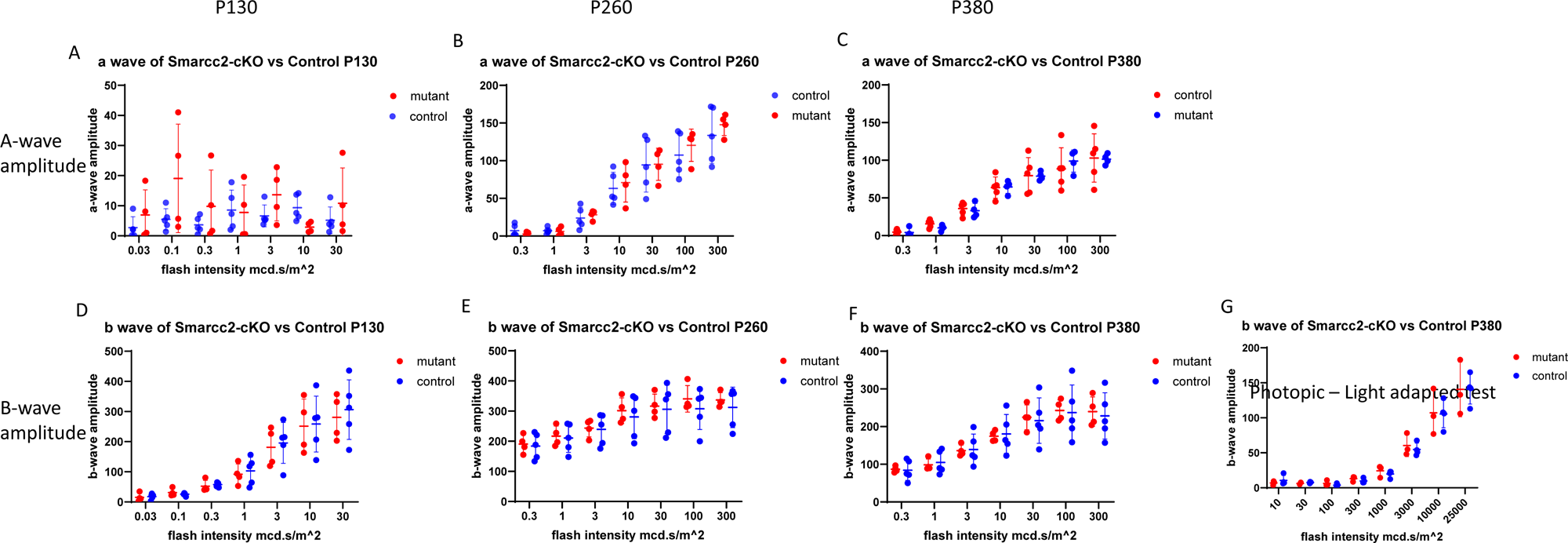
ERG test for *Smarcc2-cKO* shows no reduction in retinal response along aging compared to control. ERG tests for Control and *Smarcc2-cKO* showing amplitude of a-wave (A-C) and b-wave (D-F) of scotopic test on mice aged P130 (A, D), P260 (B, E), and P380 (C, F, n=4). b-wave of photopic ERG test on aged P380 mice (G, n=4).

**Fig. S3.**
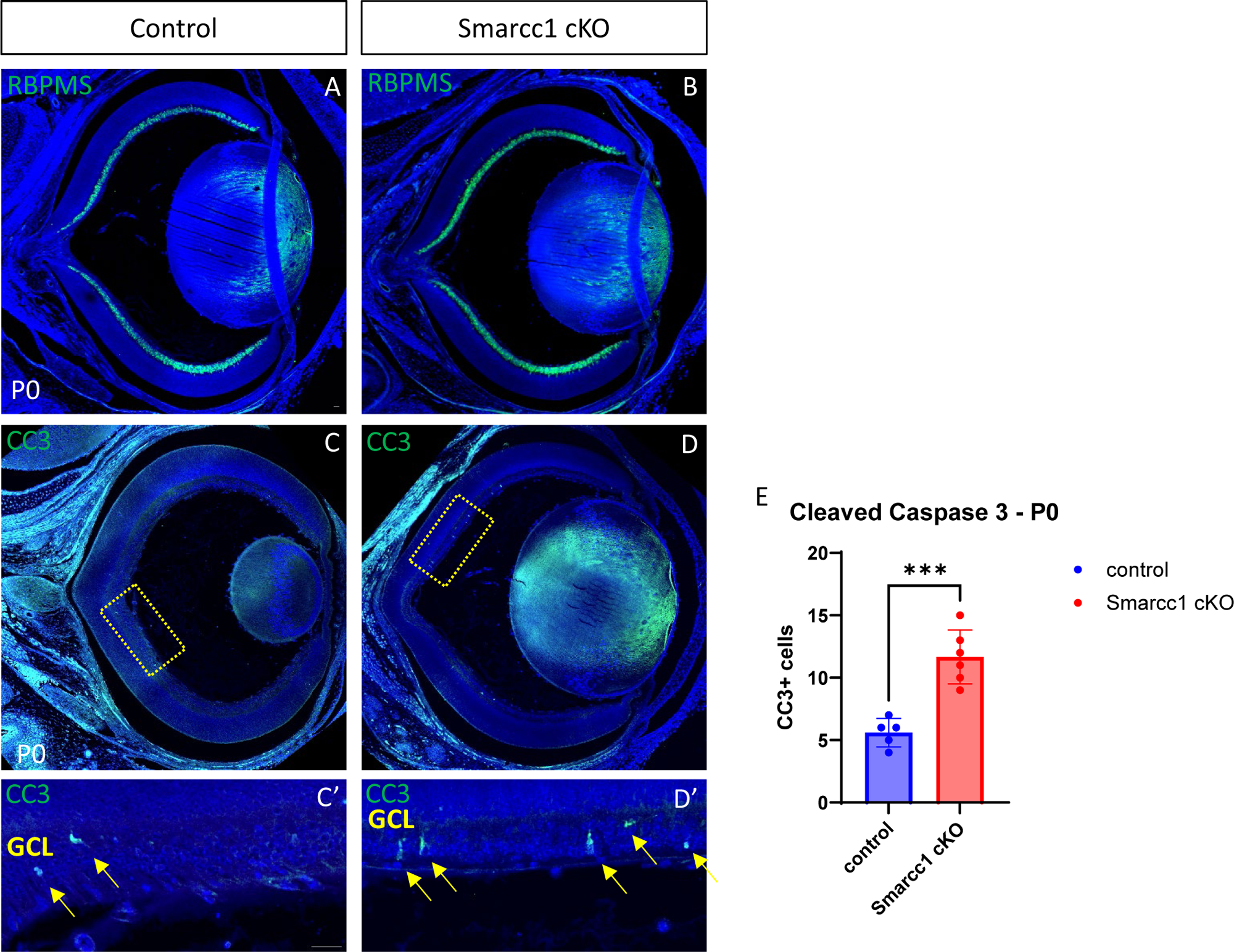
*Smarcc1-cKO* leads to increased cell death of RGCs. Antibody labeling for RGCs marker RBPMS (A-B) and apoptosis marker cleaved caspase 3 (CC3) (C-D) with respective magnifications (C’, D’) of P0 mice. Quantification of CC3 positive cells (E) in the ganglion cell layer (GCL) of control and *Smarcc1-cKO* P0 mice (n=3). Scale bars = 25µm. T-test P-val *<0.05, **<0.01, ***<0.001, ****<0.0001.

**Fig. S4.**
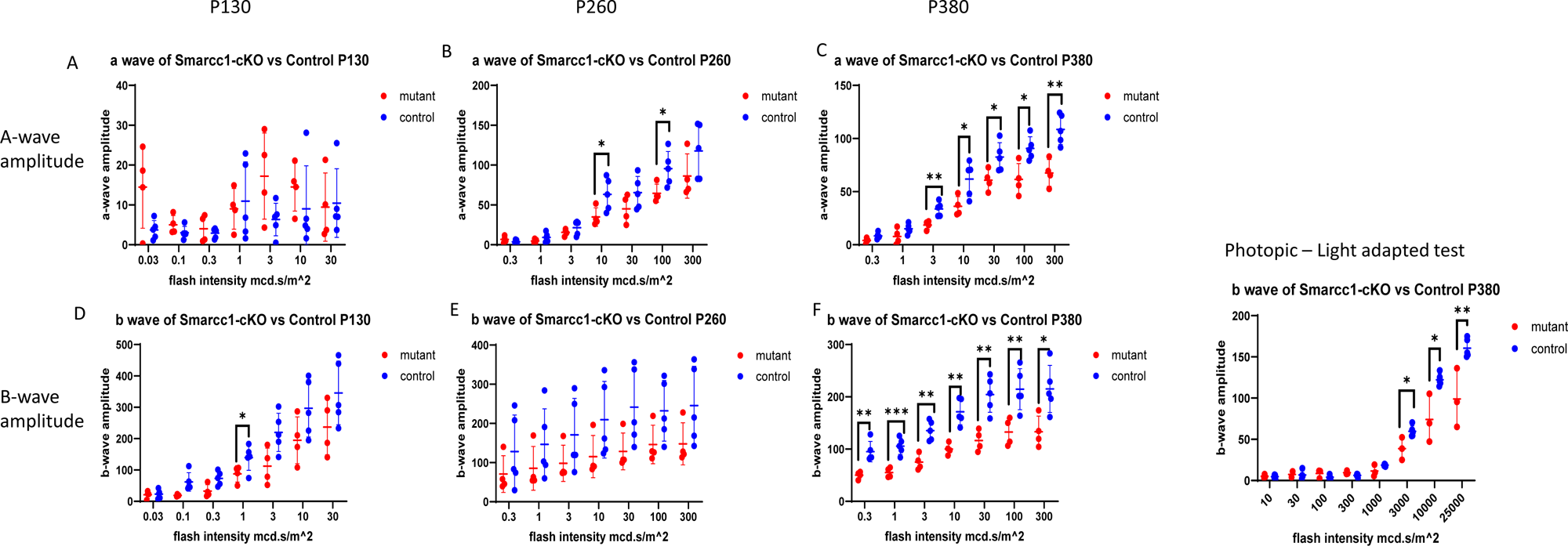
ERG test for *Smarcc1-cKO* demonstrates a reduction in retinal response with aging compared to control. ERG tests for control and *Smarcc1-cKO* display the amplitude of a-wave (A-C) and b-wave (D-F) during scotopic testing on mice aged P130 (A, D), P260 (B, E), and P380 (C, F, n=4). b-wave of photopic ERG test on aged P380 mice (G, n=4). T-test P-val *<0.05, **<0.01, ***<0.001, ****<0.0001.

**Fig. 5S.**
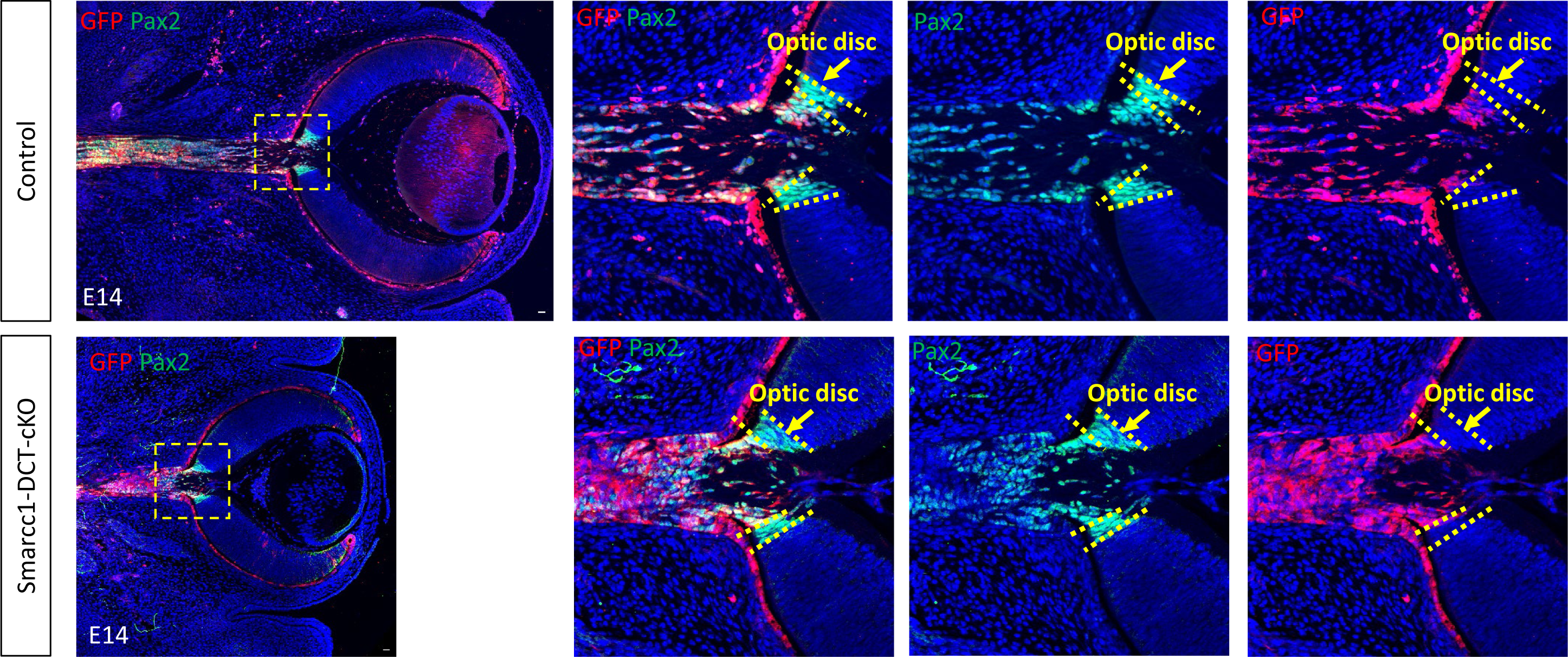
Violin plots illustrating key quality metrics of Geo-seq samples (E12, n = 3–5 per region across 10 regions), including: the number of mapped reads (in millions), mitochondrial gene percentage, mapping ratio, and the number of detected genes (TPM > 0) (A). The enrichment analysis of domain-specific genes by GSVA (B). GSEA analysis for the 3729 genes of pRPE gene set showing enrichment for genes associated light stimulus and telencephalon development (C).

**Fig. S6.**
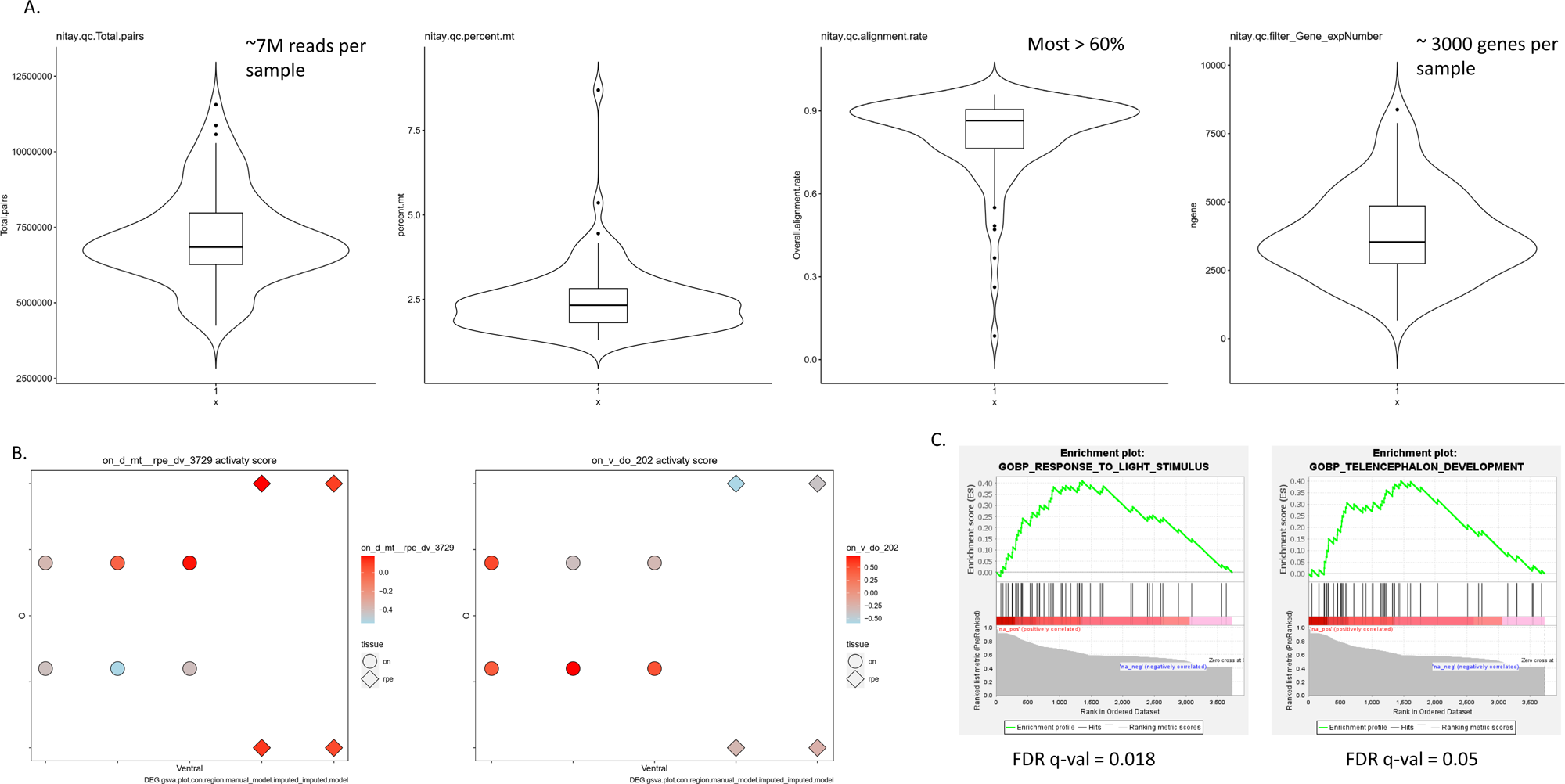
OS cross sections of E12 (A-C, G-I) and E14 (D-F, J-L) mouse embryos stained with antibodies against Smarcc1 (A,G,D,J), Smarcc2 (B,H,E,K), and Smarca4 (C,I,F,L). Scale bars = 25µm.

**Fig. S7.**
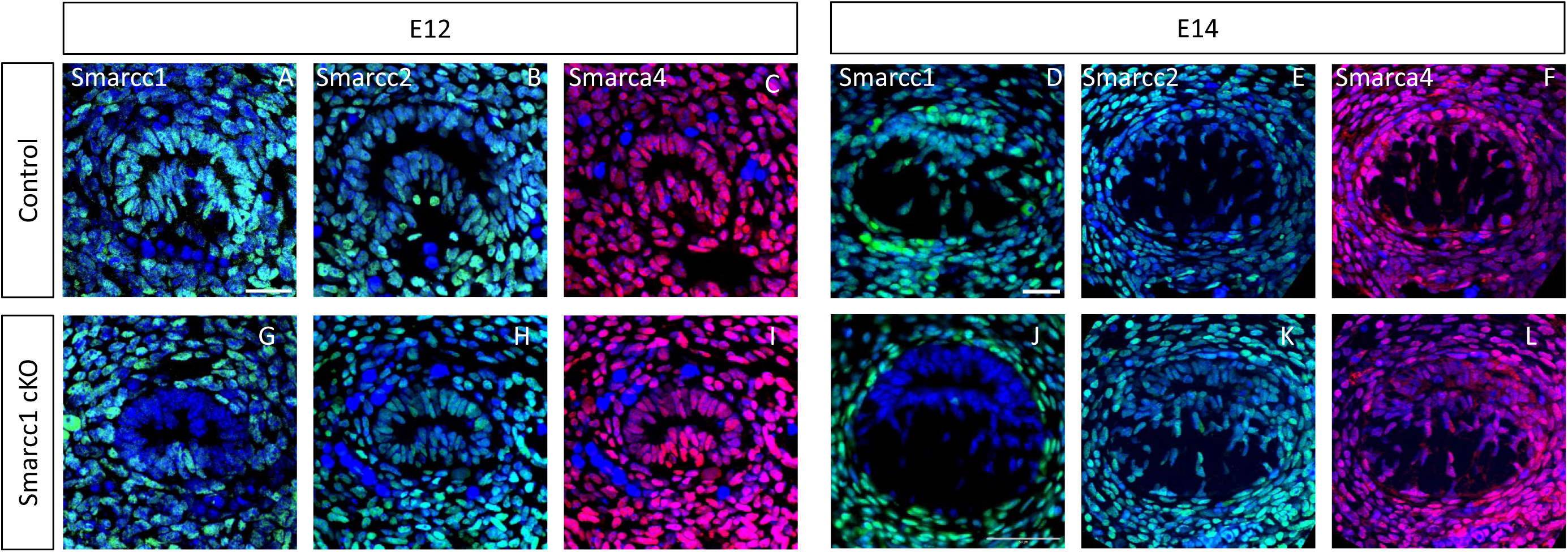
Violin plots illustrating key quality metrics of Geo-seq samples (E14, n = 3–5 per region across 3 regions), including: mitochondrial gene percentage, mapping ratio, and the number of detected genes (TPM > 0) (A).

**Fig. S8.**
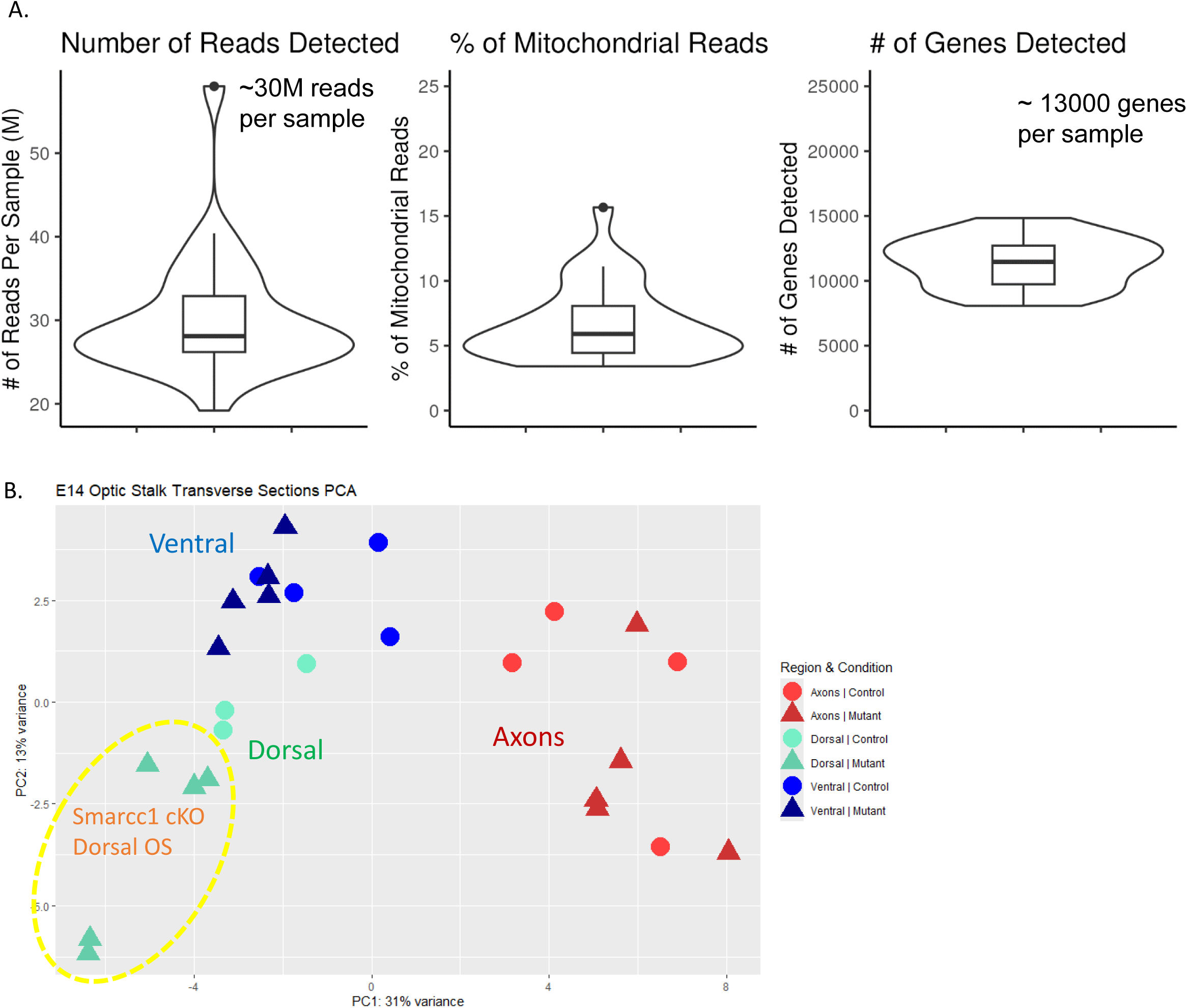
Otx2 expression during optic stalk differentiation. Antibody labeling for Otx2 in E12 (A), E13 (B) and E14 (C) mouse embryos. Higher magnifications in lower panel. Scale bars = 25µm.

**Fig. S9.**
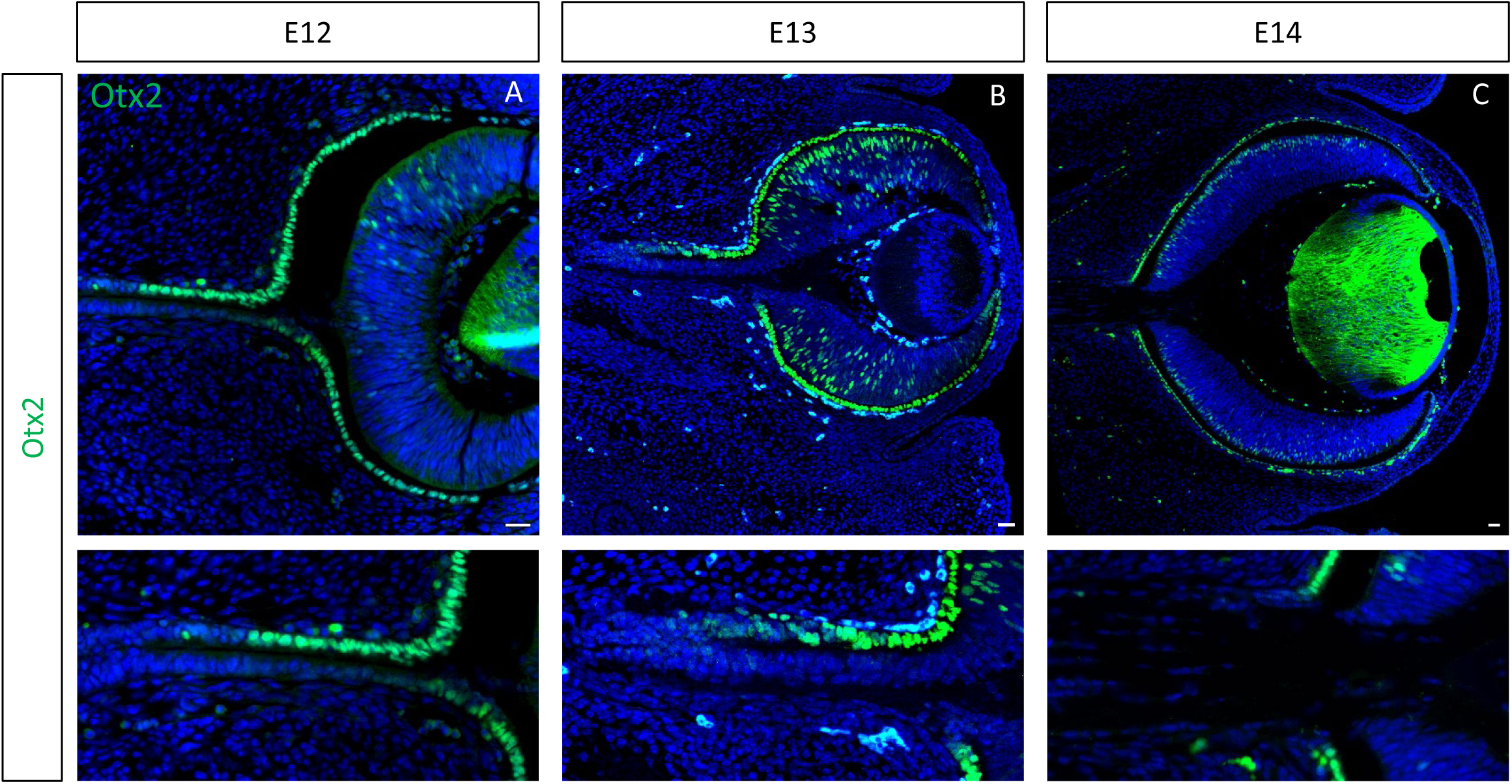
Representative samples of EdU labeling in the developing OS in control (A- D) and Smarcc1-cKO (E-H) in the dorsal and ventral optic-stalk in E12 (A, E), E13 (B, F), E14 (D, G) mouse embryos. The counts on E16 OS were conducted on the whole OS as there are no morphological differences along the dorsal ventral axes (D, H). The number of cells, based on DAPI labeling, in each region of the OS in the indicated stages (I). T-test P-val *<0.05, **<0.01, ***<0.001, ****<0.0001.

**Fig. S10.**
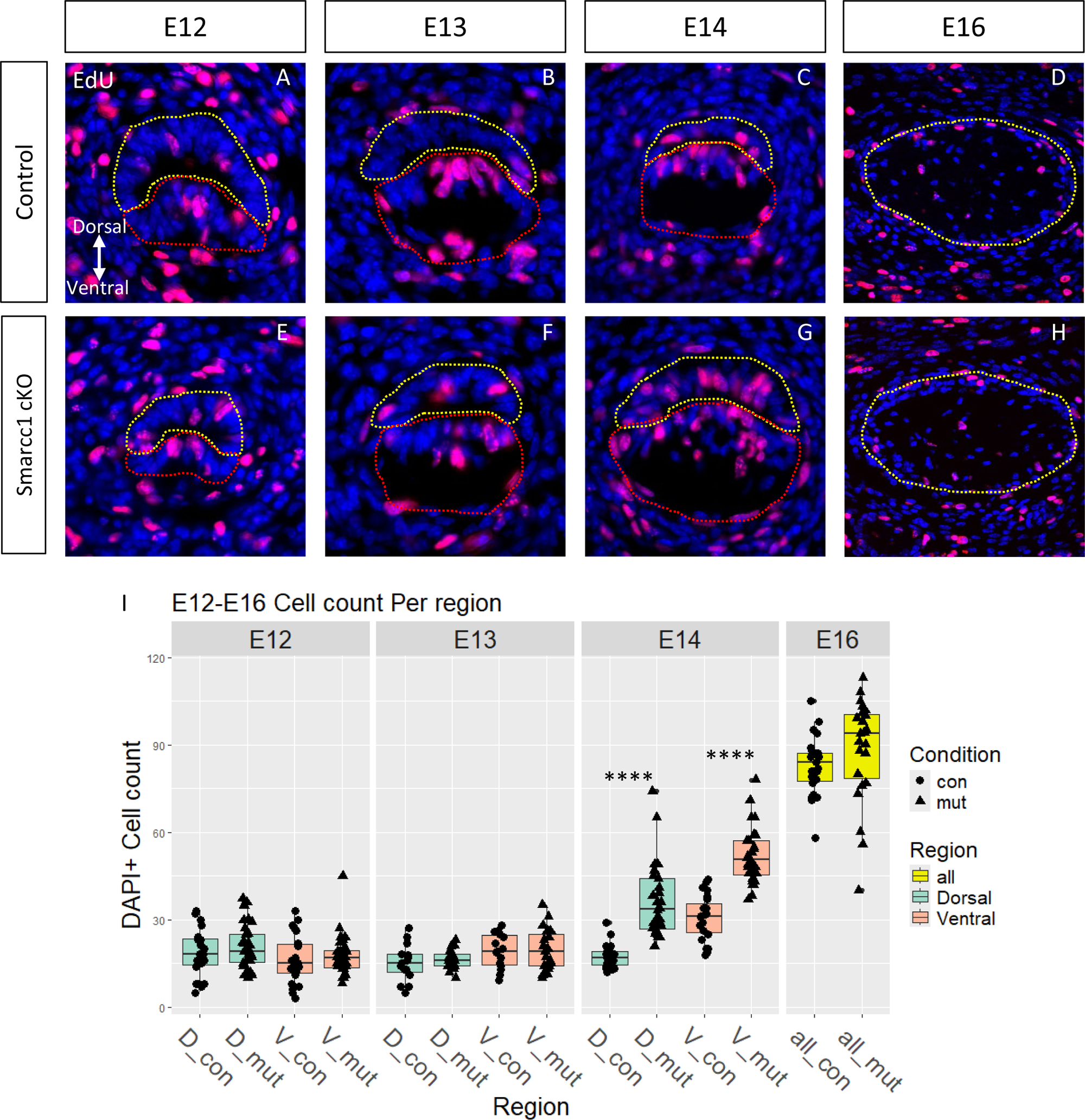
The *DCT-Cre* transgene is active in some of the optic disc progenitor cells. Antibody labeling for eGFP and Pax2 in E14 mouse embryo eye sections of control (A) and Smarcc1-DCT-cKO (B). The higher magnifications and separate channels are the right panels The overlap between the GFP and Pax2 labeling presents optic disc cells in which DCT-Cre is active while the cells that express only Pax2 (between the two dashed lines) are optic disc cells devoid of DCT-Cre activity. Scale bars = 25µm.

## Supplementary tables

Table S1_E12_LCM_Sample data and gene counts: Tab 1 – Details of the 40 E12 LCM samples. Tab 2 – First column shows gene names. The first row is sample numbers as described in the “Sample Number” column in tab 1.

Table S2_E12_Signature_Genes_Per_Region: Tab 1 – Significant genes expressed in the RPE and dorsal-distal OS domain, 3729 genes. Tab 2 – Significant genes expressed in the Ventral and dorsal-proximal OS domain, 202 genes.

Table S3_GSEA_E12: GSEA gene list for the two groups enriched in the E12 RPE and dorsal-distal OS domain. Tab 1 – GOBP_RESPONSE_TO_LIGHT_STIMULUS. Tab 2 – GOBP_TELENCEPHALON_DEVELOPMENT.

Table S4_E14_LCM_Sample data and gene counts: Tab 1 – Details of the 39 E14 LCM samples. Tab 2 – First column shows gene names. The first row is sample names as described in the “Name” column in tab 1.

Table S5_E14_DEGs_WholeGeneList: Full lists of the DEGs calculations with DESeq2 as described in Fig. 5R scheme. Tab 1 – Dorsal Control vs Smarcc1-cKO. Tab 2 – Ventral Control vs Smarcc1-cKO. Tab 3 – Axons Control vs Smarcc1-cKO. Tab 4 – Control only – Dorsal vs Ventral. Tab 5 – Control only – Dorsal vs Axons. Tab 6 – Control only – Ventral vs Axons.

Table S6_GSEA_E14: Tab 1 – GSEA gene list for the enriched groups in the E14 Smarcc1-cKO dorsal region. Tab 1 – DESCARTES_ORGANOGENESIS_NEURAL_PROGENITOR_CELLS. Tab 2 – DESCARTES_ORGANOGENESIS_MELANOCYTES. Tab 3 – RPE SIGNATURE (C1 CLUSTER) OVADIA S. ET AL. 2023.

Table S7_pcr primers table: List of PCR primers used for mouse genotyping.

Table S8_List of antibodies used: List of primary and secondary antibodies used

